# A practical modification to a resting state fMRI protocol for improved characterization of cerebrovascular function

**DOI:** 10.1101/2021.02.15.431289

**Authors:** Rachael C. Stickland, Kristina M. Zvolanek, Stefano Moia, Apoorva Ayyagari, César Caballero-Gaudes, Molly G. Bright

## Abstract

Cerebrovascular reactivity (CVR), defined here as the Blood Oxygenation Level Dependent (BOLD) response to a CO_2_ pressure change, is a useful metric of cerebrovascular function. Both the amplitude and the timing (hemodynamic lag) of the CVR response can bring insight into the nature of a cerebrovascular pathology and aid in understanding noise confounds when using functional Magnetic Resonance Imaging (fMRI) to study neural activity. This research assessed a practical modification to a typical resting-state fMRI protocol, to improve the characterization of cerebrovascular function. In 9 healthy subjects, we modelled CVR and lag in three resting-state data segments, and in data segments which added a 2–3 minute breathing task to the start of a resting-state segment. Two different breathing tasks were used to induce fluctuations in arterial CO_2_ pressure: a breath-hold task to induce hypercapnia (CO_2_ increase) and a cued deep breathing task to induce hypocapnia (CO_2_ decrease). Our analysis produced voxel-wise estimates of the amplitude (CVR) and timing (lag) of the BOLD-fMRI response to CO_2_ by systematically shifting the CO_2_ regressor in time to optimize the model fit. This optimization inherently increases grey matter CVR values and fit statistics. The inclusion of a simple breathing task, compared to a resting-state scan only, increases the number of voxels in the brain that have a significant relationship between CO_2_ and BOLD-fMRI signals, and improves our confidence in the plausibility of voxel-wise CVR and hemodynamic lag estimates. We demonstrate the clinical utility and feasibility of this protocol in an incidental finding of Moyamoya disease, and explore the possibilities and challenges of using this protocol in younger populations. This hybrid protocol has direct applications for CVR mapping in both research and clinical settings and wider applications for fMRI denoising and interpretation.

## 1. INTRODUCTION

Brain blood flow is regulated by changes in vessel diameter, directed by changes in perfusion pressure and by metabolic demands of neural activity [1]. Cerebrovascular Reactivity (CVR), the blood flow response to a vasoactive stimulus, is a metric that reflects this regulatory ability and is a key means of assessing cerebrovascular health. CO_2_ is a potent vasodilator and the partial pressure of arterial CO_2_ (PaCO_2_) naturally fluctuates with changes in respiratory depth and rate. Within a certain range around resting PaCO_2_, an increase in PaCO_2_ will cause vasodilation and a decrease will cause vasoconstriction [1]–[3]; this change in vessel diameter will result in a global change in blood flow that can be captured by any functional Magnetic Resonance Imaging (fMRI) contrast that is dependent on blood flow changes. Driven by the same physiological mechanism, the influence of PaCO_2_ on fMRI signals can either provide useful information about vascular function, or confound our measurement of neural function, depending on how one models and interprets these effects. An ideal fMRI experiment should therefore include characterization of CVR, both to provide complementary vascular information and to better model and interpret any neural activity of interest (e.g., task activation, intrinsic fluctuations, and functional connectivity). The main focus of this paper is to assess how a practical modification of a typical “resting-state” protocol improves CVR mapping, focusing on regional variations in semi-quantitative CVR amplitude and local hemodynamic timings. Considering broader applications, improved modeling of PaCO_2_ fluctuations in any fMRI data naturally enables better differentiation between non-neuronal confounds and neuronally-driven effects and can aid in correcting fMRI analyses for variations in transit delays and vascular properties of the local hemodynamic response [4]–[10].

### 1.1. Practical CVR mapping: modelling both amplitude and timing

Here we used the Blood Oxygenation Level Dependent (BOLD) fMRI response to represent blood flow changes, and the partial pressure of end tidal CO_2_ (P_ET_CO_2_) to represent PaCO_2_ changes [11], [12]. Aside from an invasive contrast-agent such as acetazolamide, CO_2_ gas inhalation methods are often seen as the gold standard for CVR mapping with fMRI [13]. Gas inhalations allow more precise and repeatable PaCO_2_ targeting, however these experiments are more timely, costly and complicated to set-up, and are therefore not practical for all research and clinical applications. Much previous fMRI research has demonstrated that CVR mapping with breathing tasks (breath holding, BH, or cued deep breathing, CDB) is a promising practical approach that can provide useful information about cerebrovascular health [14], [15]; this has been demonstrated in a diverse set of clinical cohorts, e.g. [16]–[30]. Resting P_ET_CO_2_ fluctuations also have a significant positive relationship with BOLD fMRI signals [31]. Therefore, an even simpler approach to CVR mapping is to measure natural fluctuations in P_ET_CO_2_ during fMRI acquisitions with no specific task, i.e., during rest [15], [32], [33]. This may be favored in clinical studies where subject compliance with breathing tasks is hard to achieve. The utility of this resting-state CVR approach has also been demonstrated in clinical cohorts [32], [34]–[37].

Both breathing tasks and resting-state approaches produce comparable BOLD signal changes [9], [38], [39], and are also comparable to those obtained with gas-inhalation techniques [9], [32], [33], [40], [41]. There are few studies comparing breathing tasks and resting-fluctuations for CVR mapping normalized to a common scale, i.e., P_ET_CO_2_. One study, using P_ET_CO_2_ regressors in their CVR analysis, report that resting-state data shows poorer model fits, poorer repeatability, and more variable between-subject CVR estimates compared to BH data, in 14 subjects [42]. CVR maps showed good spatial agreement between BH CVR and resting-state CVR when the latter is evaluated based on the resting P_ET_CO_2_ trace and the resting state fluctuation amplitude (RSFA), but in general their results suggest it is not straight-forward to replace BH designs with resting-state in the assessment of CVR. In terms of agreement in CVR timing, [43] reported a strong agreement between P_ET_CO_2_ latency values derived from a BH dataset and resting-state dataset, in one subject. Also assessing timing, [44] investigated the optimal temporal shift between a grey matter (GM) BOLD time-series and a P_ET_CO_2_ regressor, in 12 subjects, within a 16 second range. In the modelled resting-state data, some subjects showed negative correlation values, and no clear shift maximum within the temporal bounds considered. For the modelled BH data, all subjects showed a significant positive correlation between BOLD and P_ET_CO_2_ that peaks at a physiologically plausible temporal shift. Further, BH derived optimal shift values were repeatable within two halves of the scan, whereas this was not the case for optimal shift values derived with resting-state data. Though CVR mapping with resting-state data is possible, there exists intrinsic low-frequency oscillations, driven by neural activity or other physiological processes [45]–[49] that can be of similar or greater magnitude to the low-frequency fluctuations induced by P_ET_CO_2_, sometimes resulting in an fMRI time-course poorly coupled to P_ET_CO_2_. Furthermore, breathing tasks induce larger fluctuations in P_ET_CO_2_ and therefore larger fMRI signal changes which can be easier to detect above noise. However, breathing tasks, as opposed to rest, can introduce motion confounds that are correlated with task timings [50]–[52].

Correcting for the temporal offset between a P_ET_CO_2_ regressor and the local fMRI response is an important and necessary step in estimating accurate regional CVR values. Though the previous literature has mixed approaches and results, robustly characterizing this temporal shift in resting-state data sometimes appears unreliable and less repeatable. Within the BOLD fMRI literature, it appears relatively common to correct for the temporal offset with a cross-correlation between the physiological regressor and an average fMRI regressor. It is less common to model this temporal offset on a voxel-wise basis, though there are multiple examples in the literature showing the implementation and advantages of this in resting-state or breathing task data [6], [29], [43], [50], [53]– [62]. This temporal offset is driven by both methodological and physiological factors: there is a delay between the CO_2_ exhalation inside the scanner and the recording of exhaled CO_2_ in the control room, vascular transit delays as gases travel with the blood to arrive at each brain region and variability in the vasodilatory response of local arterioles and the spatio-temporal complexities of the BOLD response. Therefore, it is important to model CVR lag (also referred to as CVR timing, optimal shift, temporal offset, latency or delay) on a regional or voxel-wise basis. In healthy subjects, we recently demonstrated our approach to voxel-wise optimization of hemodynamic lag, to improve regional BOLD-CVR estimates [63], [64], and we apply this pipeline to our CVR mapping analysis in this paper. As well as improving model fit and more accurately characterizing CVR amplitude, making maps of hemodynamic lag can provide distinct regional information that is clinically relevant [61], [65], [66] and potentially aid in correcting fMRI analyses [4]–[10].

There is always a trade-off between complexity of experimental set-up and how much control one can have over the manipulation of blood gases. We strive for simple and feasible methods that can be applied in clinical settings, without losing too much accuracy, and minimizing the disruption to the overall scan session. Therefore, in this study we propose a practical addition to a typical resting-state fMRI scan: approximately 2.5 minutes of a breathing task appended to the start of a resting state period. We suggest that this novel hybrid design (breathing task + resting state) will be useful for both mapping of CVR amplitude and timings, ideally still allowing for separate analysis of resting state data. We compare CVR maps with and without lag optimization, and CVR maps that have been created with resting-state data alone, resting-state data preceded by a short hypercapnic breathing task, and resting-state data preceded by a short hypocapnic breathing task. We chose two different breathing tasks to achieve these PaCO_2_ changes: the commonly utilized breath-hold (BH) task to induce hypercapnia (reviewed by [14]), and a cued deep breathing task (CDB) to induce hypocapnia via hyperventilation [50], [58], [67], both tasks reviewed by [15].

## 2. METHODS

### 2.1. Data Collection

This study was reviewed by Northwestern University’s Institutional Review Board and all subjects gave written informed consent. Nine healthy subjects were recruited (6 female, mean age = 26.22±4.06 years). A tenth subject was recruited of which a potential incidental finding was observed, based on hemodynamic lag maps. The appropriate ethical guidelines were followed in reporting of this incidental finding, and it was later confirmed this subject had a diagnosis of Moyamoya disease. Therefore, this subject is not described alongside the other nine subjects in this manuscript, but as a case study in a separate section of the results.

The overall study design is shown in Figure 1. Before scanning, subjects practiced the BH and CDB tasks outside the scanner with the researcher (R.C.S). Three fMRI scans were collected, the order of acquisition pseudo-randomized across subjects (Figure 1). BH and CDB timings were guided by previous work using these tasks [42], [50] including information about the BOLD time to peak and return to baseline timings in response to a single deep breath [57]. From three fMRI scans, five data segments were created: BH + REST, CDB + REST, REST, REST_BH_ and REST_CDB_, of which the first two contain breathing tasks and the others do not. Each data segment had the same number of time points, to match degrees of freedom across models. During scanning, inspired and expired CO_2_ and O_2_ pressures (in mmHg) were sampled through a nasal cannula worn by the participant, and pulse was monitored with a finger transducer. Although pulse data were collected, they were not included in the modeling of fMRI data due to insufficient quality of the recordings across many subjects and scans. During the fMRI scans, volume triggers were recorded. All signals were recorded at 1000Hz with LabChart software (v8.1.13, ADInstruments), connected to a ML206 Gas Analyzer and PL3508 PowerLab 8/35 (ADInstruments).

**Figure 1.**
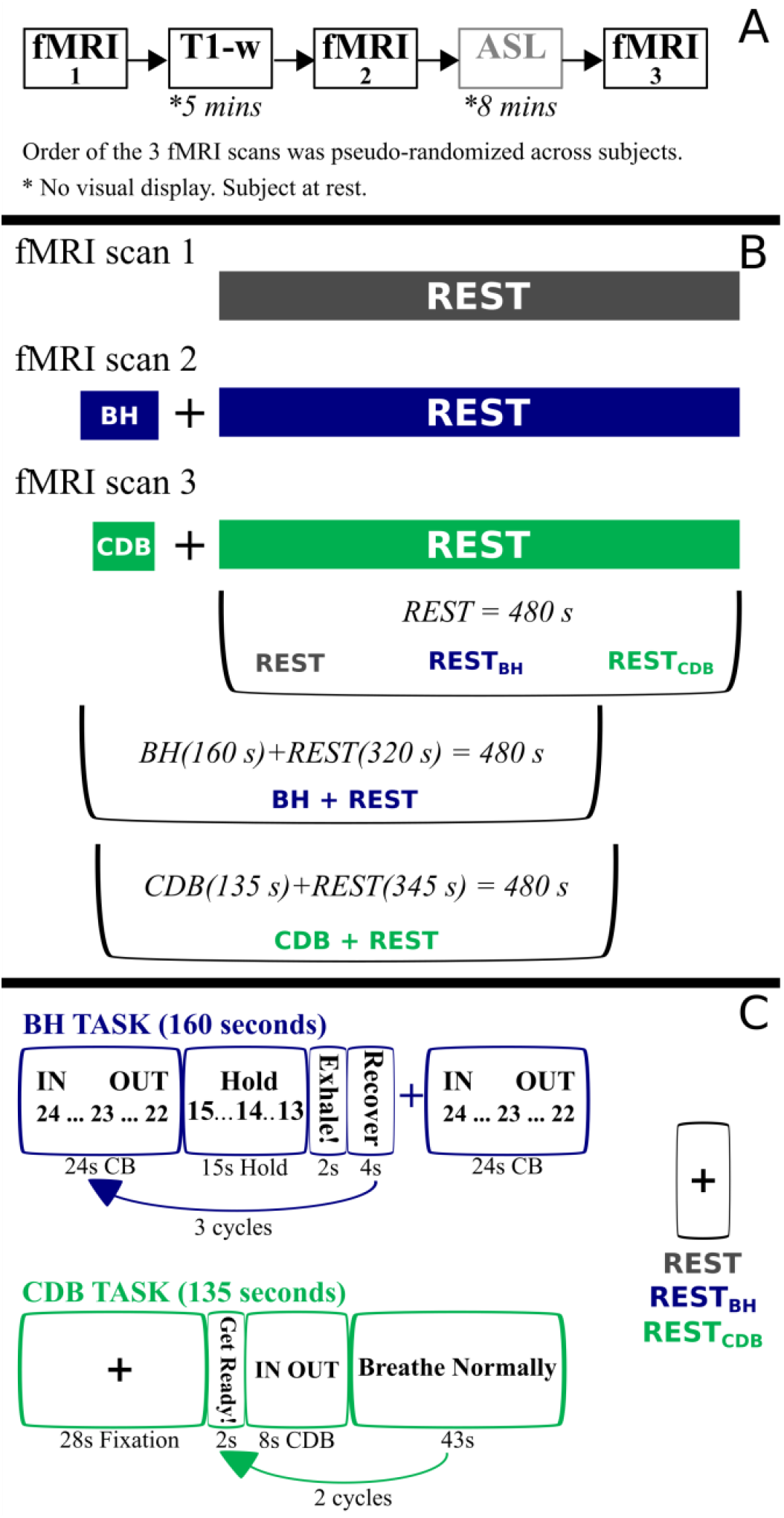
(A) Five scans collected during the whole session (40 minutes). The ASL scan is not analyzed in this manuscript. (B) From 3 fMRI scans, 5 data segments were extracted, each with the same number of time points: 3 segments were REST only and the other 2 segments involved a breathing task (BH/CDB) followed by REST. Visual instructions for each task were displayed on a monitor during scanning. (C) BH task: IN and OUT instructions alternated for 3 s each, with a countdown from 24 s. Subjects ended on an exhale before holding, and were instructed to do another exhale after holding. ‘Recover’ is a period of free breathing. CDB task: IN and OUT instructions alternated for 2 s and subjects were told to take fast, deep breaths. REST: fixation cross shown. BH = breath holding, CDB = cued deep breathing, CB = cued breathing.

Imaging data were collected with a Siemens 3T Prisma MRI system with a 64-channel head coil. The functional T2*-weighted acquisitions were gradient-echo planar sequences provided by the Center for Magnetic Resonance Research (CMRR, Minnesota) with the following parameters: TR/TE = 1200/34.4 ms, FA = 62°, Multi-Band (MB) acceleration factor = 4, 60 axial slices with an ascending interleaved order, 2 mm isotropic voxels, FOV = 208 x 208 mm^2^, Phase Encoding = AP, phase partial Fourier = 7/8, Bandwidth = 2290 Hz/Px. Single-band reference (SBRef) images were also acquired to facilitate functional realignment and masking. A whole brain T1-weighted EPI-navigated multi-echo MPRAGE scan was acquired, adapted from [68], with these parameters: 1 mm isotropic resolution, 176 sagittal slices, TR/TE1/TE2/TE3 = 2170/1.69/3.55/5.41 ms, TI = 1160 ms, FA = 7°, FOV = 256×256, Bandwidth = 650 Hz, acquisition time of 5 minutes 12 seconds, including 24 reacquisition TRs. The three echo images were combined using root-mean-square. ASL data was also collected before the last fMRI scan, but it was not analyzed in the current study.

Five example datasets from a pediatric study of hemiparetic cerebral palsy and typical development (ages 7-21 years, all female) are included to assess the feasibility of our proposed method in cohorts where task compliance may be more challenging. All gave written informed consent or assent. Only one functional T2*-weighted acquisition was collected. The functional acquisition matched the parameters explained previously, except for these key differences: MB factor = 8, TR/TE=555/22 ms, FA = 47°, 64 slices, 6/8 phase partial Fourier, and FOV = 208×192. During this acquisition participants completed a CDB+REST protocol that matched the timings described previously, except auditory cues were used instead of visual cues (Figure 10). Expired CO_2_ was collected as previously described.

### 2.2. Data Analysis

The data from this study unfortunately cannot be made openly available due to restrictions of the ethical approval that they were collected under. However, analysis derivatives that are not included in this manuscript may be provided, on request, within ethical guidelines. All breathing task stimulus code and the main analysis code have been made available via this GitHub repository: github.com/BrightLab-ANVIL/Stickland_2021

#### 2.2.1. MRI pre-processing

A custom shell script grouped AFNI [69] and FSL [70]–[73] commands, for minimal preprocessing of the MRI data. DICOMS were converted to NIFTI format with dcm2niiX [72]. The T1-weighted file was processed with FSL’s *fsl_anat*, involving brain extraction [74], bias field correction and tissue segmentation (GM/white matter/cerebral spinal fluid) with FAST [75]. Tissue masks were subsequently created by thresholding the partial volume estimate images at 0.75. The Single Band Reference image (SBRef) from the middle (second) fMRI scan was brain extracted, and eroded. The SBRef image was registered to the preprocessed T1-weighted image using FLIRT [76], [77]. The transformation matrix was inverted in order to co-register the tissue masks from T1 image space to SBRef image space. For each fMRI acquisition, the first 10 volumes were discarded to allow the signal to achieve a steady state of magnetization. AFNI’s *3dvolreg* was run with the same middle SBRef scan as the reference volume. Six motion parameters (three translations, three rotations) were saved and demeaned for each acquisition. Next, the three fMRI files were masked to brain voxels using the SBRef mask created previously.

#### 2.2.2. CO_2_ trace pre-processing

Custom MATLAB (MathWorks, R2018b) code processed the physiological recording to create P_ET_CO_2_ regressors. A text file from the whole scan protocol was exported from the LabChart software, and this text file was split into an output for each fMRI acquisition. Each output was (purposefully) slightly longer than the length of the fMRI acquisition: an additional 20.4 seconds of data (equivalent to 17 extra TRs) both before and after each acquisition was included in the exported data. This made it possible to create shifted P_ET_CO_2_ regressors in a later step. The output for each functional acquisition was processed with a peak detection algorithm to detect the end-tidal peaks (maximum CO_2_ value at the end of each exhale). The output of the peak detection algorithm was manually checked for every dataset to ensure the end of each expiration breath was always chosen, and to remove incorrectly identified end-tidal values (e.g., in the case of partial breathing through the mouth, and not fully through the nose, not giving a true end-tidal peak). A linear interpolation between these peaks produced the P_ET_CO_2_ trace (for the breath-hold periods in which there is no end-tidal recording, a linear interpolation is based on the last exhale before the hold and the first exhale after), which was then convolved with the SPM canonical hemodynamic response function (HRF). The resultant P_ET_CO_2_hrf was exported for functional imaging analyses.

#### 2.2.3. P_ET_CO_2_ regressors at different temporal shifts

101 shifted P_ET_CO_2_hrf regressors of the same length were created, with different temporal offsets. Starting with P_ET_CO_2_hrf regressor timings that overlapped with the fMRI acquisition, the regressor was shifted back and shifted forward in time, in 0.3 second increments. We recently presented this approach for hemodynamic lag optimization [63], however, in that earlier implementation we included an additional step prior to performing this ‘fine shift’ (shifting in 0.3 s to create 101 regressors): a cross-correlation between the P_ET_CO_2_hrf trace and the up-sampled GM time-course as an initial gross re-alignment (‘bulk shift’) to account for measurement delay and a general vascular transit delay. In our previous work, which involved breath-hold data only, the fine shifted regressors were shifted *relative* to the bulk shifted regressor, up to a shift maximum of ±9 s, based on what had been seen in previous literature. In this current work, we chose not to bulk shift via the cross-correlation method before running the lagged generalized linear model (GLM) analysis, as some bulk shifts for the REST data segments were found to be very far from physiological reasonable values (e.g., 19 and 20 seconds at the most extreme, see **Figure 4**), and deviating greatly from the optimum shift found in BH+REST or CDB+REST data segments. We hypothesize that this could be due to the intrinsic low-frequency oscillations associated with neural activity that can be of similar or greater magnitude to the low-frequency fluctuations induced by P_ET_CO_2_, resulting in a GM fMRI time-course poorly coupled to P_ET_CO_2_. Motion and other physiological noise could also contribute. Therefore, bulk shifting in these datasets seemed inappropriate, as errors would be produced in the early stages of our analysis pipeline, likely biasing the resultant comparisons of lag optimized CVR maps across data segments. Instead, we chose to apply our voxel-wise lag optimization method with a larger shift maximum, searching for the optimum shift (i.e., the hemodynamic lag) within ±15 s from the original P_ET_CO_2_hrf regressor. This 15 second range was based on a 9 second range plus the largest bulk shift seen across subjects for the BH+REST and CDB+REST data segments, which was −5.45 seconds. A range of ±9 s was justified previously [63], and is consistent with much research in healthy subjects reporting lag values or breathing task regressors within this range around an average value [6], [43], [50], [57]–[59], [61], [78].

#### 2.2.4. CVR and lag estimation

Figure 2 shows the mean BOLD time-series across GM voxels for each fMRI acquisition and each subject, and also the P_ET_CO_2_hrf (unshifted) time-series for reference. CVR modelling was carried out separately for each of the five data segments illustrated in Figure 1 and 2.

**Figure 2.**
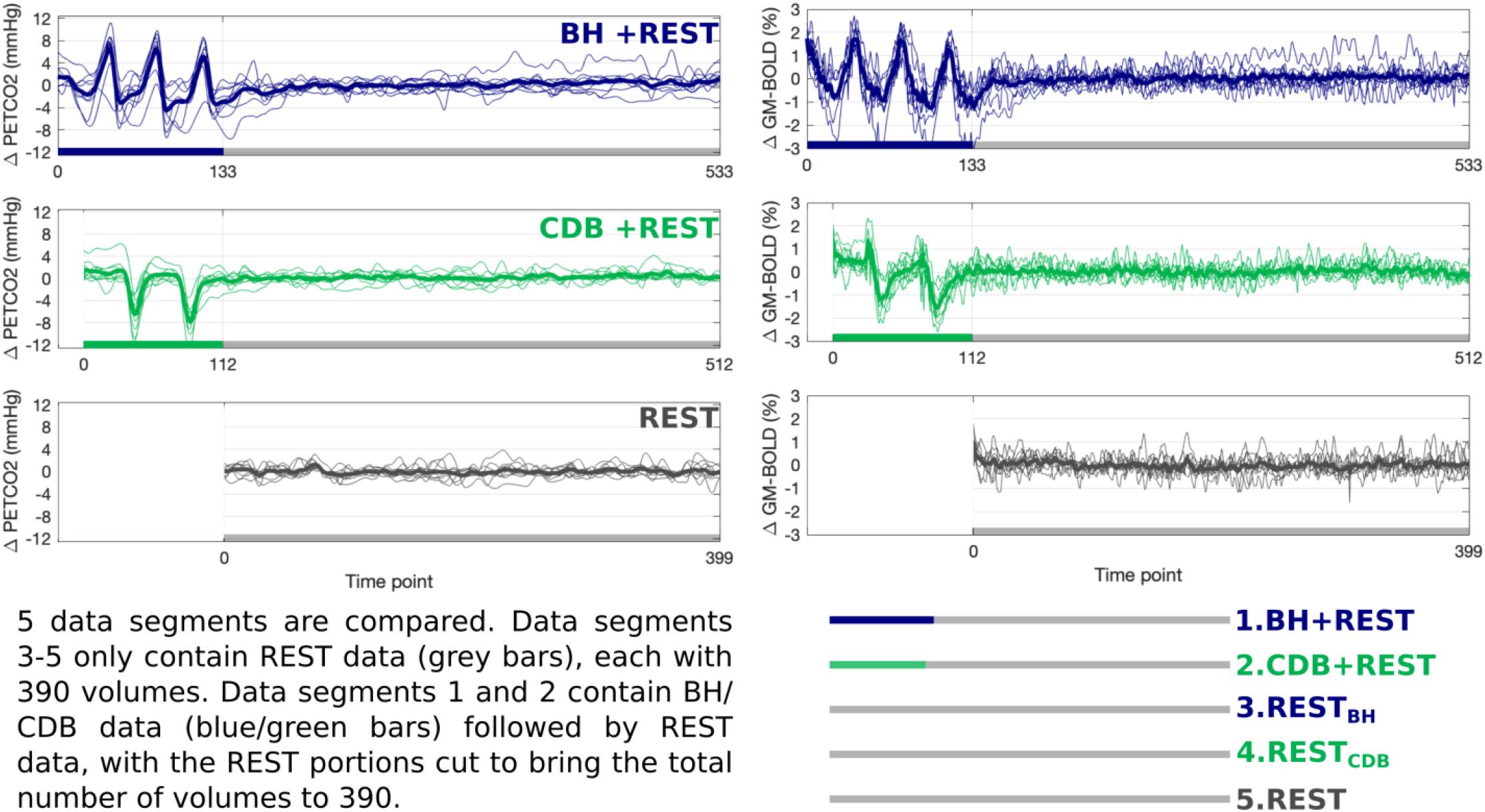
The left column displays unshifted P_ET_CO_2_hrf traces (mmHg change from baseline) and the right column displays GM-BOLD traces (% change from mean) for each of three fMRI acquisitions. Thick lines represent group means, and thin lines represent each subject. The key at the bottom describes the five data segments that are compared in this manuscript. The first 10 volumes at the start of each data segment are not used (discarded to allow steady-state to be reached) resulting in 390 volumes for each data segment.

**Figure 3.**
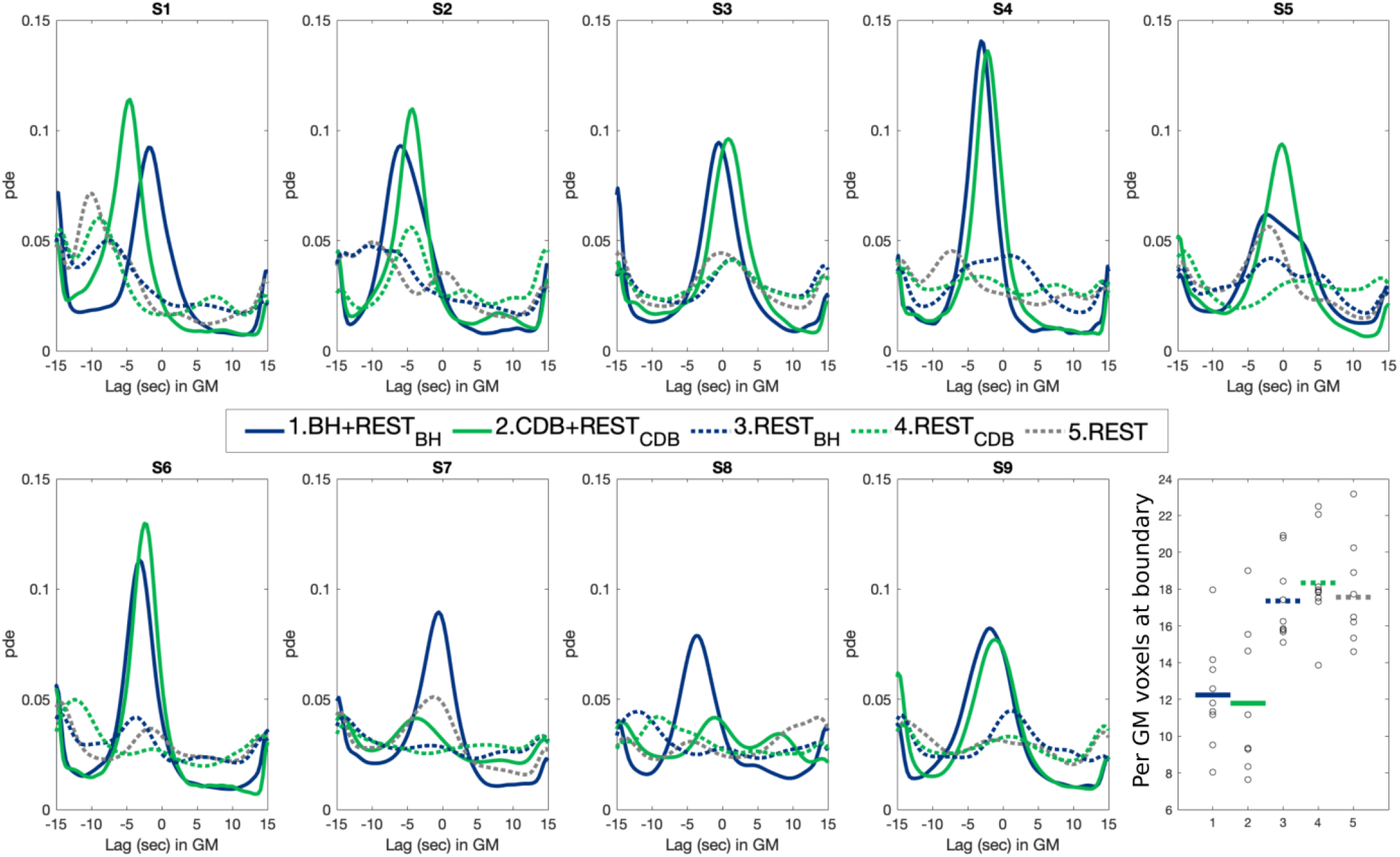
Distributions of lag values, across GM voxels, expressed as probability density estimates (pde). Distributions for all subjects (S1-S9) are shown, and for all data segments. The percentage (per) of GM voxels with optimal shift values at the boundary condition is compared across data segments in the bottom right-hand corner. The group mean is shown as a thick horizontal line and each subject is represented as a black dot.

**Figure 4.**
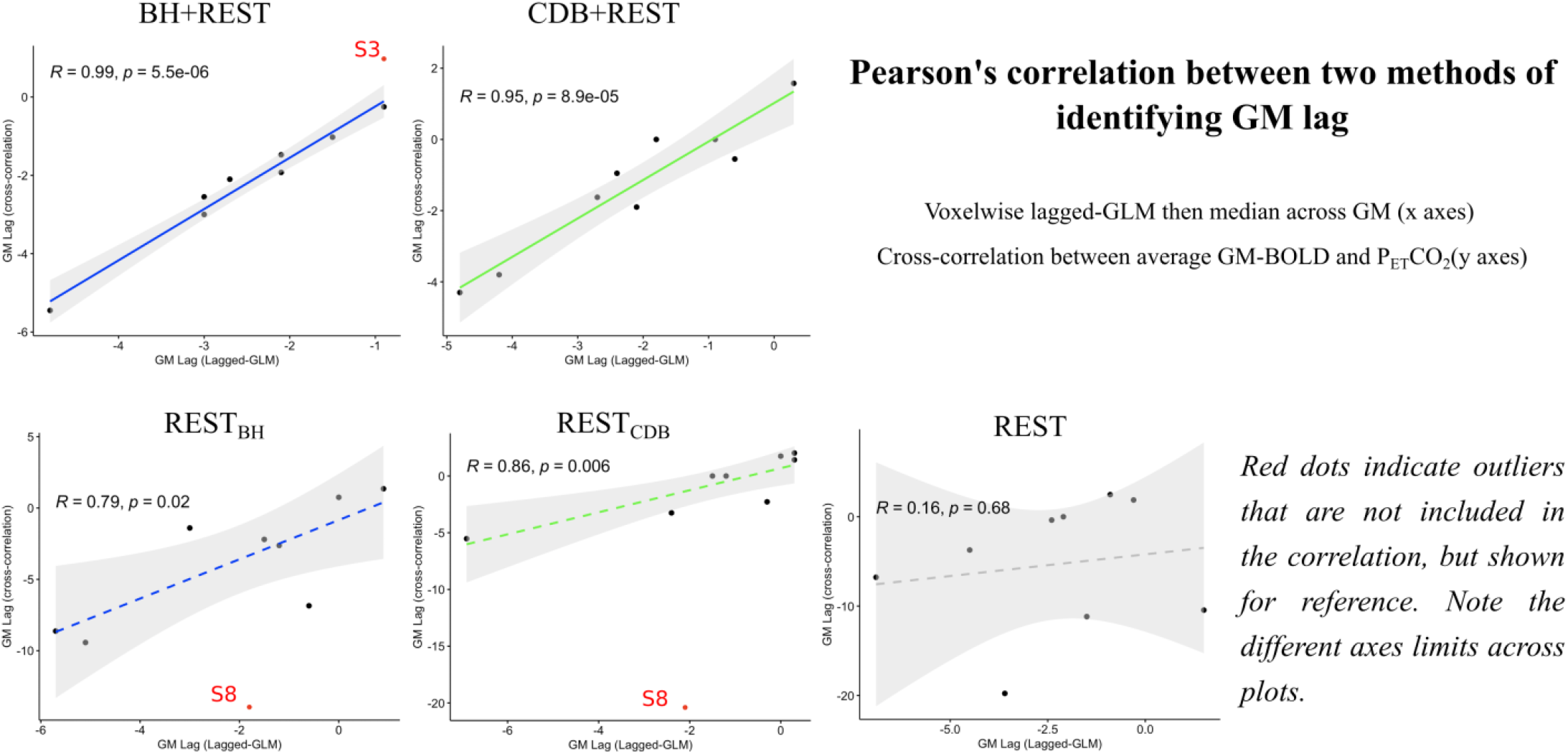
Correlation between the median lag over GM voxels from the voxelwise lagged-GLM analysis and a cross-correlation between the average GM-BOLD time-series and the P_ET_CO_2_hrf time-series. Each dot represents one subject, and correlations are shown for each data segment. The grey shaded regions around the fit line indicate the 95% confidence interval of the correlation coefficient. Data points were classed as outliers (indicated by red dots) when a Cook’s distance was over 4/n, with n being the number of subjects. P-values are FDR corrected.

A custom shell script grouped AFNI and MATLAB (Version 2018b, MathWorks) commands to create maps of CVR and hemodynamic lag. The fMRI variance explained by the demeaned P_ET_CO_2_hrf regressor was modeled alongside the six demeaned motion parameters (as nuisance regressors) and Legendre polynomials up to the 4th degree (to model the mean and drifts of the fMRI signal). Including these polynomials in the model is approximately equivalent to a high-pass filter with a cutoff of 0.0076Hz (AFNI 3dDeconvolve help; [79]). Least squares regression accounting for serial autocorrelation of residuals was applied with AFNI’s *3dREMLfit* command. The beta coefficient for the P_ET_CO_2_hrf was scaled by the fitted mean of that voxel to create CVR maps in %BOLD/mmHg. This same model was run for all shifted versions of the P_ET_CO_2_hrf regressor. This lagged GLM approach is explained in our previous work [63]: the hemodynamic lag at each voxel was identified as the shift that gave the maximum full model coefficient of determination (R^2^). Final hemodynamic lag values ranging from negative to positive indicate earlier to later hemodynamic responses, respectively. As a result, two CVR maps were created: with no lag optimization (No-Opt) and CVR maps with lag optimization (Lag-Opt). Lag-Opt CVR maps used the beta coefficient for the P_ET_CO_2_hrf regressor from the model with the optimum shift. If a voxel with an optimal shift (lag) was found at or adjacent to a boundary (−15, −14.6, +14.6, +15) this was not deemed a true optimization and is also less likely to be physiologically plausible. These voxels are not included when plotting or summarizing lag and Lag-Opt CVR maps. Voxel-wise CVR values were deemed significant for absolute T-statistics greater than 1.96, corresponding to p<0.05. For Lag-Opt CVR, this threshold was further adjusted with the Šidák correction [44], [80] due to the lagged approach running 101 different GLMs.

#### 2.2.5. Data summaries and statistical tests

The median GM CVR and the percentage of significant voxels in GM was calculated for each modelled data segment. These values were computed for No-Opt, Lag-Opt, No-Opt with statistical thresholding (p<0.05), Lag-Opt with matched statistical thresholding (p<0.05) and Lag-Opt with stricter statistical thresholding (p<0.05, Šidák corrected). The kernel density estimation of the distribution of lag values in GM (MATLAB’s *ksdensity* function) was also computed for each subject and each data segment, and the median GM lag values were outputted.

In order to provide lag values with some regional specificity across GM regions, FSL atlases in MNI space (MNI-maxprob-thr25-2mm and HarvardOxford-sub-maxprob-thr25-2mm) were used to make three GM masks: cortical GM, subcortical GM and cerebellar GM. From the HarvardOxford atlas, left and right cerebral cortex parcels were combined into one mask, and left and right subcortical regions (thalamus, caudate, putamen, palladium, hippocampus, amygdala, accumbens) combined into another. The cerebellum parcel was extracted from the MNI atlas to make a third mask. These three atlas masks (cortical, subcortical and cerebellar) were linearly transformed (FSL, FLIRT) to subject space, and thresholded to only include voxels within the subject’s GM tissue mask. Median lag values were extracted from each mask for BH+REST and CDB+REST data segments.

R version 3.4.1 (R Core Team, 2019) was used for data exploration and statistical testing. To compare parameter values across data segments and optimization schemes (No-Opt vs Lag-Opt), repeated measures ANOVAs were run with the R package permuco [82] with the ‘aovperm’ function. Null distributions were created via 100,000 permutations of the original data, which therefore do not depend on gaussian and sphericity assumptions. When investigating simple main effects (‘emmeans’ package) and performing multiple comparisons, p-values were adjusted with the Benjamini & Hochberg approach [83] for control of false discovery rate (FDR), and then compared against an alpha of 0.05 to determine significance. Outliers were identified with boxplots. Correlation plots and statistical outputs were created with the R packages ggplot2 [84] and ggpubr [85] with the ‘ggscatter’ function. Outliers for the correlation analysis were identified when Cook’s distance was over 4/n, with n being the number of subjects, which indicates an influential single data point. The Shapiro-Wilk test was used to ensure normality of variables.

It can be seen from the parameter maps (Figure 5, Figure 6, Supplementary Figure 1) that after statistical thresholding, a substantial proportion of voxels within white matter do not show a significant relationship between P_ET_CO_2_hrf and BOLD signals. Therefore, we decided to focus our comparison of CVR and lag parameter averages, across the different data segments, within GM voxels only.

**Figure 5.**
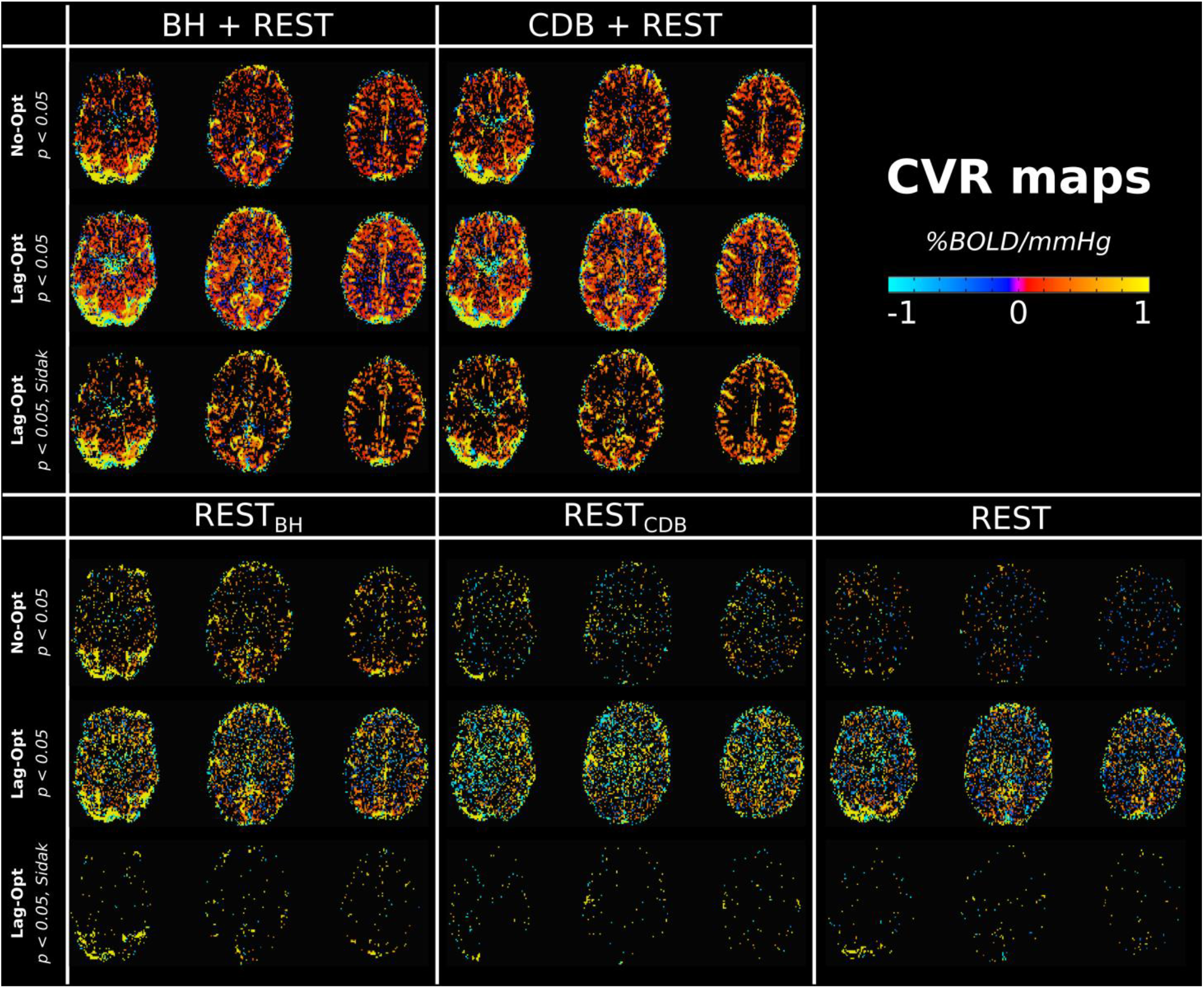
Maps of Cerebrovascular Reactivity (CVR) for one example subject (S4), displayed for each of the five data segments. For each data segment, CVR maps are shown with no lag optimization (No-Opt) thresholded with the P_ET_CO_2_hrf regressor at p<0.05 (top row for each data segment), with lag optimization (Lag-Opt) thresholded with p<0.05 but no Sidak correction (middle row) and thresholded with p<0.05 with Sidak correction (bottom row). CVR (Lag-Opt) maps do not include voxels with optimum lags found at the boundary.

**Figure 6.**
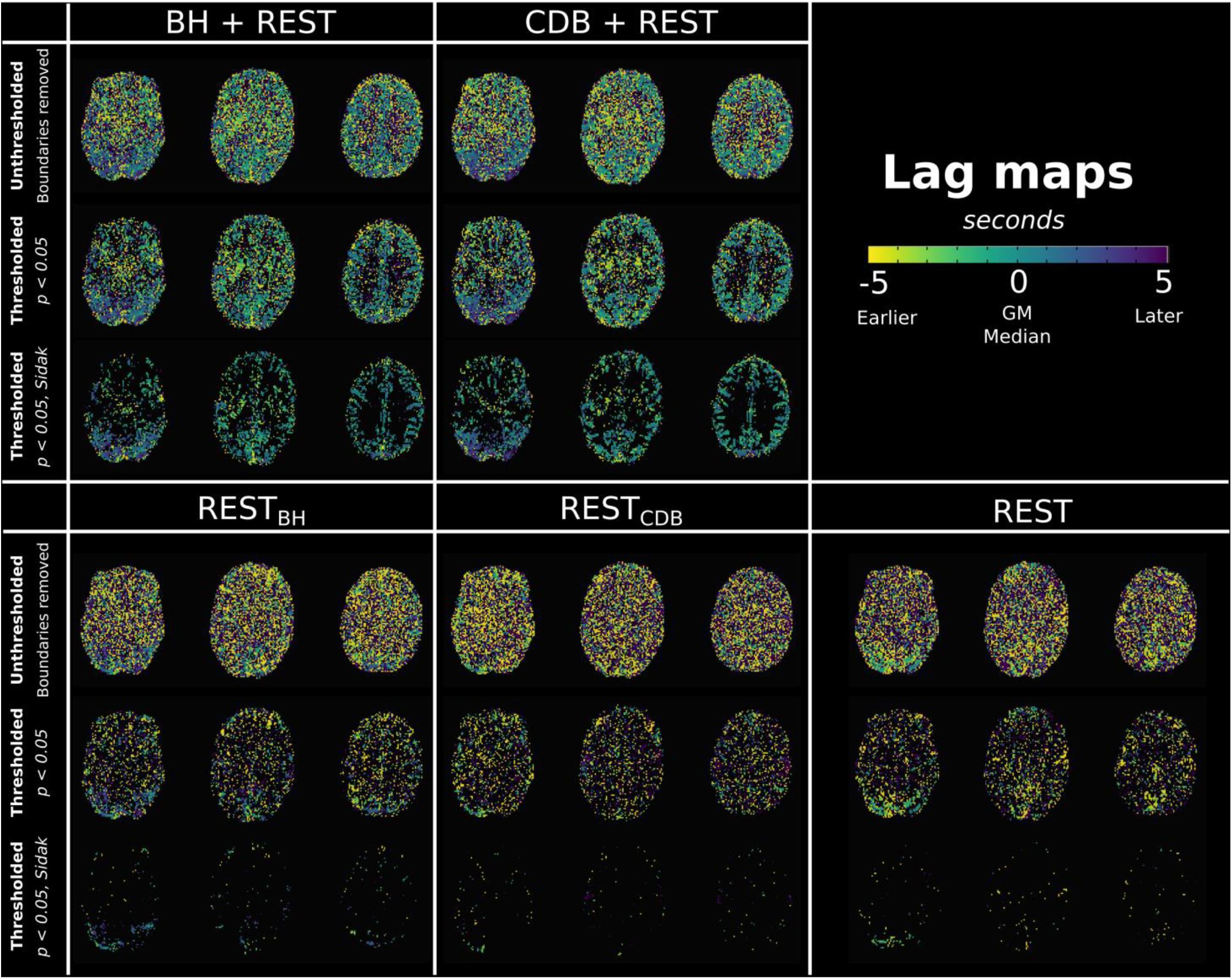
Maps of hemodynamic lag for one example subject (S4), displayed for each of the five data segments. For each data segment, lag maps are shown without statistical threshold (top row for each data segment), thresholded with the P_ET_CO_2_hrf regressor at p<0.05 and no Sidak correction (middle row), and thresholded with p<0.05 with Sidak correction (bottom row). All lag maps do not include voxels with optimum lags found at the boundary.

## 3. RESULTS

Figure 2 shows the average BOLD-fMRI signal across GM, and the corresponding P_ET_CO_2_hrf changes, for each of the three scans. Note the three cycles of increased P_ET_CO_2_ values in blue, due to breath holds, and the two cycles of decreased P_ET_CO_2_ values in green, due to cued deep breaths. The fMRI time-series changes as expected due to the CO_2_ manipulations. The BH task produced a maximum pressure increase of 7.4±3.1 mmHg (mean±stdev across subjects) and CDB produced a maximum decrease of 8.1±3.2 mmHg, relative to the mean value at rest. For the three rest segments the temporal standard deviation of P_ET_CO_2_ was 0.9±0.4, 1.2±0.7 and 0.8±0.4 mmHg (mean±stdev across subjects) for REST, REST_BH_ and REST_CDB_ respectively.

### 3.1. Hemodynamic lag values

Figure 3 shows the distribution of hemodynamic lag values in GM tissue estimated from the lagged-GLM analysis, displaying both positive and negative lags. Though a positive lag value might be expected (CO_2_ pressure change in the blood *leads* the fMRI signal change) there are multiple factors contributing to this value, and in different directions. The recording delay between CO_2_ exhalation inside the scanner and CO_2_ recording outside the scanner contributes to a more negative shift, whereas vascular transit delays and vasodilatory dynamics that eventually lead to the BOLD-fMRI signal will contribute to a more positive shift. Therefore, the observed lag between a CO_2_ recording and a BOLD fMRI recording will naturally vary with the experimental set-up, the participant, and their physiological state. Due to this, the lag parameter maps displayed and summarized later in this manuscript are normalized to be relative to the GM median value. Such normalization allows more valid comparisons when summarizing across subjects and comparing with other literature, showing the spatial variability of hemodynamic lag.

As demonstrated in Figure 3, the BH+REST and CDB+REST data segments, plotted in solid blue and green lines respectively, have a more gaussian-like distribution of lag values (excluding the boundaries), the properties of which generally match for most subjects. REST data segments (dashed lines) have less physiologically plausible distributions which agree less with other segments and vary more across subjects. Figure 3 also shows the percentage of GM voxels with lag values at the boundary condition. From the repeated measures ANOVA analysis, there was a significant effect of data segment on percentage of voxels at the boundary condition (F(4,32) = 9.86, p < 0.0001). Simple main effects analysis showed that group means for BH+REST (12.24%) and CDB+REST (11.79%) did not differ (p=0.795, FDR-corrected). However, BH+REST and CDB+REST each had a significantly lower percentage compared to REST (17.56%), REST_BH_ (17.35%) and REST_CDB_ (18.34%), with all p-values < 0.022, FDR corrected. All REST segment pairs were not significantly different to each other (p>0.605, FDR corrected). This analysis was run multiple times after the separate removal of three data points from the REST_CDB_ group which were classed as extreme outliers based on boxplots, however the results did not change. The three REST data segments resulted in a greater percentage of voxels being identified at the boundaries of our fitting procedure; lag values at the boundary are less physiologically plausible, and indicate less certain lag optimization i.e., we cannot determine this is a true local maximum.

We compared the GM median lag value, obtained with this voxel-wise lagged-GLM analysis, with a lag obtained when performing a cross-correlation between the P_ET_CO_2_hrf time-series and the mean BOLD fMRI time-series across all GM (as is commonly done in the literature, when no voxel-wise correction is applied, described as ‘bulk shift’ in the methods). This evaluation is shown in Figure 4 for each data segment, where each point in each plot corresponds to one subject. Across these two different methods of characterizing a representative GM lag value, the BH+REST and CDB+REST data segments show the strongest significant positive correlations. The REST_BH_ and REST_CDB_ segments also show strong significant positive correlations, though the range of values for the cross-correlation method is much greater than for the lagged-GLM method, and there is one extreme outlier. The REST segment did not show a significant correlation between methods. These results indicate that there is clear consistency between different methods of summarizing a GM lag value when including either breathing task; this consistency is present in some, but not all, REST segments. Considering the shape of the lag distributions in Figure 3 it is important to acknowledge that the use of the median as a summary metric may not be completely valid for all subjects and data segments.

Supplementary Table 1 shows that BOLD response timing to a P_ET_CO_2_hrf change is generally earliest in subcortical GM regions, approximately 0.4 seconds later in cortical GM regions, and 1.4–2.0 seconds later in GM cerebellar regions. Though specific regional comparisons were not a focus of this paper, lag (CVR delay) is less commonly characterized in the literature, compared to CVR amplitude. Therefore, these results are included in order to assess the agreement of our lag values with previous literature.

### 3.2. CVR and lag maps

Figures 5 and 6 show maps of CVR and lag, respectively, for one subject and all thresholding options. CVR values increase after lag optimization, as expected. There is more spatial agreement in CVR and lag maps when the BH and CDB segments are included in the modelled data, showing a similar contrast between tissues types. Maps that only include REST data barely exhibit a physiologically reasonable contrast between tissues types, and after the final statistical thresholding (p<0.05, Šidák corrected) very few voxels remain. This is also seen in Supplementary Figure 1, which displays maps for all subjects. All subjects follow the trend described, except S7 and S8 which have CDB+REST maps that appear more similar to REST maps (in number and distribution of significant voxels); these are the same two subjects in Figure 3 that had CDB+REST GM lag distributions that did not look similar to the BH+REST distributions. Subject 8 was also an outlier in Figure 4, albeit for REST_BH_ and REST_CDB_.

### 3.3. Comparing GM CVR values and significant fits across data segments

Figure 7 depicts the distribution of CVR values in GM, for No-Opt CVR and Lag-Opt CVR across different levels of statistical thresholding, for one subject. The same plots can be found for all subjects in Supplementary Figures 2-6. In general, the No-Opt CVR values (row 1) follow a Gaussian- or Laplacian-like distribution when no statistical thresholding is applied. The REST data segments generally have the greatest number of negative CVR values. The Lag-Opt CVR distributions (unthresholded, row 2) change shape to resemble a more bimodal distribution, due to CVR estimates diverging further from zero in either direction (which is expected due to our method for optimizing lag). This effect is enhanced in the thresholded distributions for both No-Opt and Lag-Opt (rows 3-5); CVR values closer to zero are removed after applying statistical criteria, resulting in distinct bimodal distributions. Interestingly, we see the proportion of negative CVR values to positive CVR values is much less in the BH+REST and CDB+REST data segments. For the REST segments, after lag optimization and statistically thresholding there are some cases where there is an equivalent amount of positive and negative CVR values. We expect predominantly positive CVR values in GM, and true negative CVR responses represent very different physiological mechanisms [86], [87]. Considering the shape of these distributions, and the considerable proportion of negative CVR values in the REST segments, we summarized positive and negative CVR separately when computing summary CVR metrics across GM, shown in Figure 8 and Supplementary Figure 7. For ease of reference, group averages and standard deviations from these figures are also provided in Supplementary Table 2.

**Figure 7.**
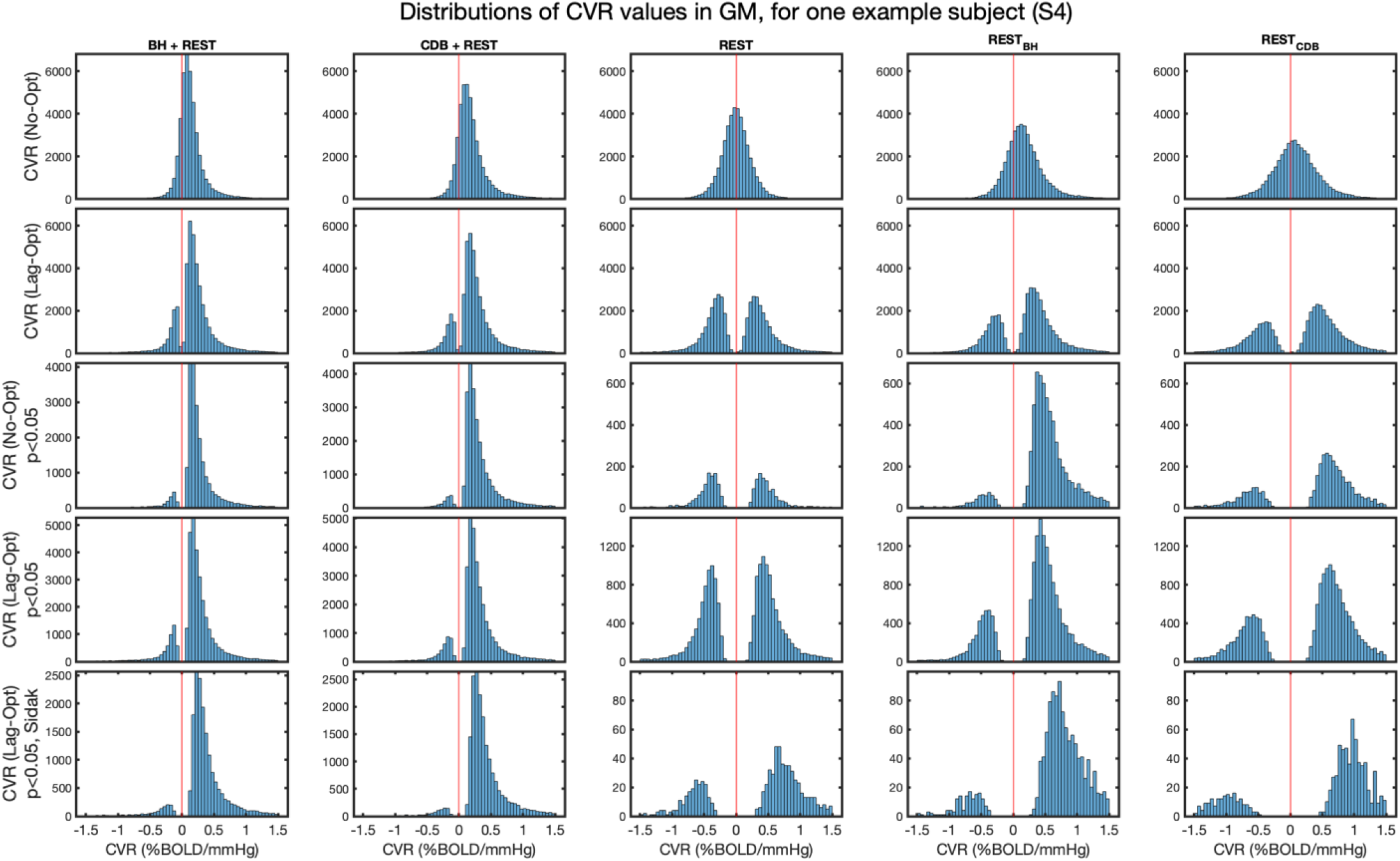
Distributions of CVR values in GM for one example subject (S4). The y-axes show the frequency count. The same scaling is used for rows 1 and 2 because no thresholding is applied and therefore all GM voxels are included. When statistical thresholding is applied (rows 3, 4 and 5), different numbers of GM voxels remain for each data segment.

**Figure 8.**
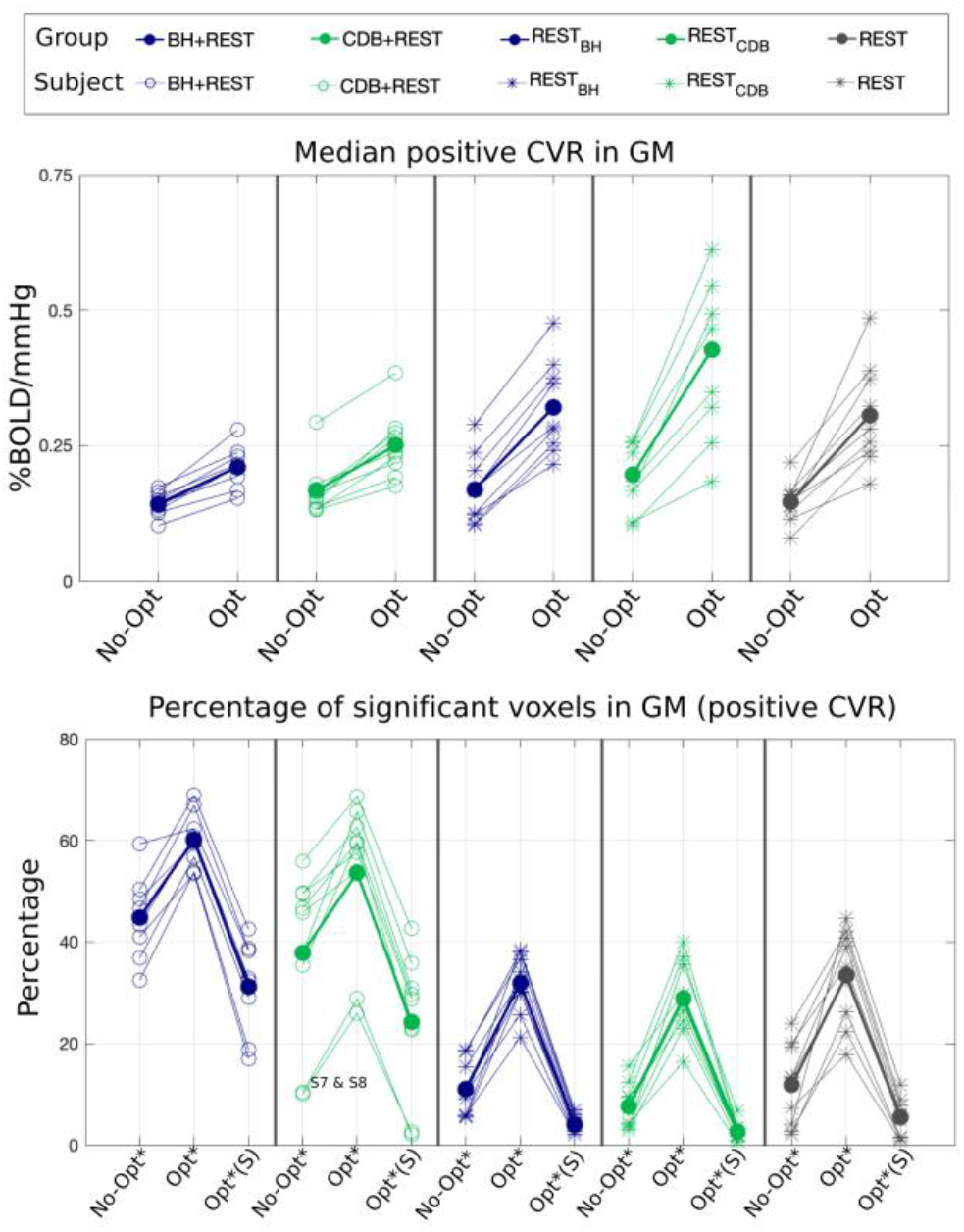
Comparing GM summary metrics across data segments. The legend corresponds to the column structure in each panel; both single subject data and group means are shown. Top: Median positive CVR across all GM voxels for non-optimized (No-Opt) and lag optimized (Opt) analyses. Bottom: Percentage of significant positive fits in GM for each data segment and level of thresholding: the * indicates thresholding at p<0.05 and the *(S) indicates thresholding at p<0.05 with Sidak correction.

Figure 8 (top panel) compares median positive GM CVR values across data segments. There was a significant interaction between data segment and optimization scheme (F(4,32 = 14.2, p<0.00001) so simple effects were investigated. Positive GM CVR values increased after lag optimization for all data segments: BH+REST (p=0.02), CDB+REST (p=0.01), REST (p=0.002), REST_BH_ (p=0.002) and REST_CDB_ (p=0.002). There were no significant differences in median positive GM CVR values between any pair of data segments for No-Opt values (all p-values >0.14) but there were significant differences for Lag-Opt values, which drove the significant interaction. For Lag-Opt, positive GM CVR values were not different between: BH+REST and CDB+REST; CDB+REST and REST; CDB and REST_BH_; REST and REST_BH_ (all p-values >0.14), However, REST had significantly higher values than BH+REST (p=0.02); REST_BH_ had significantly higher values than BH+REST (p=0.006); REST_CDB_ had significantly higher values than BH+REST (p=0.001), CDB+REST (p=0.001), REST (p=0.002), and REST_BH_ (p=0.008). One extreme outlier was removed from the CDB+REST segment (Figure 8, subject with highest values) for this statistical testing. All p-values are FDR corrected. Similar patterns are seen for negative CVR values (Supplementary Figure 7).

The previous CVR comparisons are taken using all GM voxels, and not only from voxels that are statistically thresholded; Figure 8 (bottom panel) shows that the percentage of significant voxels in GM is noticeably lower in the REST data segments, suggesting that there is less confidence in these summary CVR estimates. With or without lag optimization, there are more GM voxels showing significant positive fits for P_ET_CO_2_hrf in the data segments with breathing tasks, for all subjects except S7 and S8. The inverted V-shaped pattern shows the number of significant voxels changes with statistical thresholding in a similar way across the 5 data segments: more voxels are significant after lag optimization if the same thresholding (p<0.05) is applied, however after Šidák correction this returns to a similar or smaller number of statistically significant voxels as found without lag optimization. This shows the statistical consequence of the lagged-GLM computation (considered further in the discussion).

### 3.4. Clinical Utility

An incidental finding was suspected in one of our participants due to an abnormality first noticed in the lag maps. Specifically, a large section of the cortex displayed a blood flow response to P_ET_CO_2_ much later than other areas of the cortex. The area of cortex impacted appeared to be the vascular territory mostly supplied by the middle cerebral artery. The appropriate ethical procedures were followed and it was confirmed that this subject had Moyamoya disease. Moyamoya is a rare vascular disorder which typically involves blockage or narrowing of the carotid artery, reducing blood flow to the brain. Figure 9 shows CVR and lag maps in this subject. The lag maps show that many GM voxels in the right hemisphere are responding approximately 10 seconds later compared to the homologous regions of the left hemisphere. Note, the analysis run for this subject was the same as previously presented, however the lag values that are included in these maps are not relative to GM median, due to the clear bilateral vascular pathology, and this case being interpreted separately from the other subjects. When lag is not considered, the CVR maps (No-Opt) show dominant negative CVR in this vascular territory, similar to what has been reported in previous CVR mapping studies with Moyamoya [88]–[90]. However, when correcting the CVR maps for lag (Lag-Opt) there is a striking change -the CVR within GM mostly normalizes, without a clear pathology. These lag optimized maps suggest that the local CVR response is preserved in GM, albeit with delayed blood transits. Looking at the lag optimized CVR map alone, one could incorrectly conclude that this subject does not have a clear vascular pathology. Both maps, lag and CVR (Lag-Opt), are needed for the most accurate interpretation, and to determine whether there are CVR reductions, delayed blood transits, or both. Figure 9 also shows CVR and lag results in REST data. Here, the pathology is much less evident, and these results follow the same pattern seen in the other 9 subjects presented.

**Figure 9.**
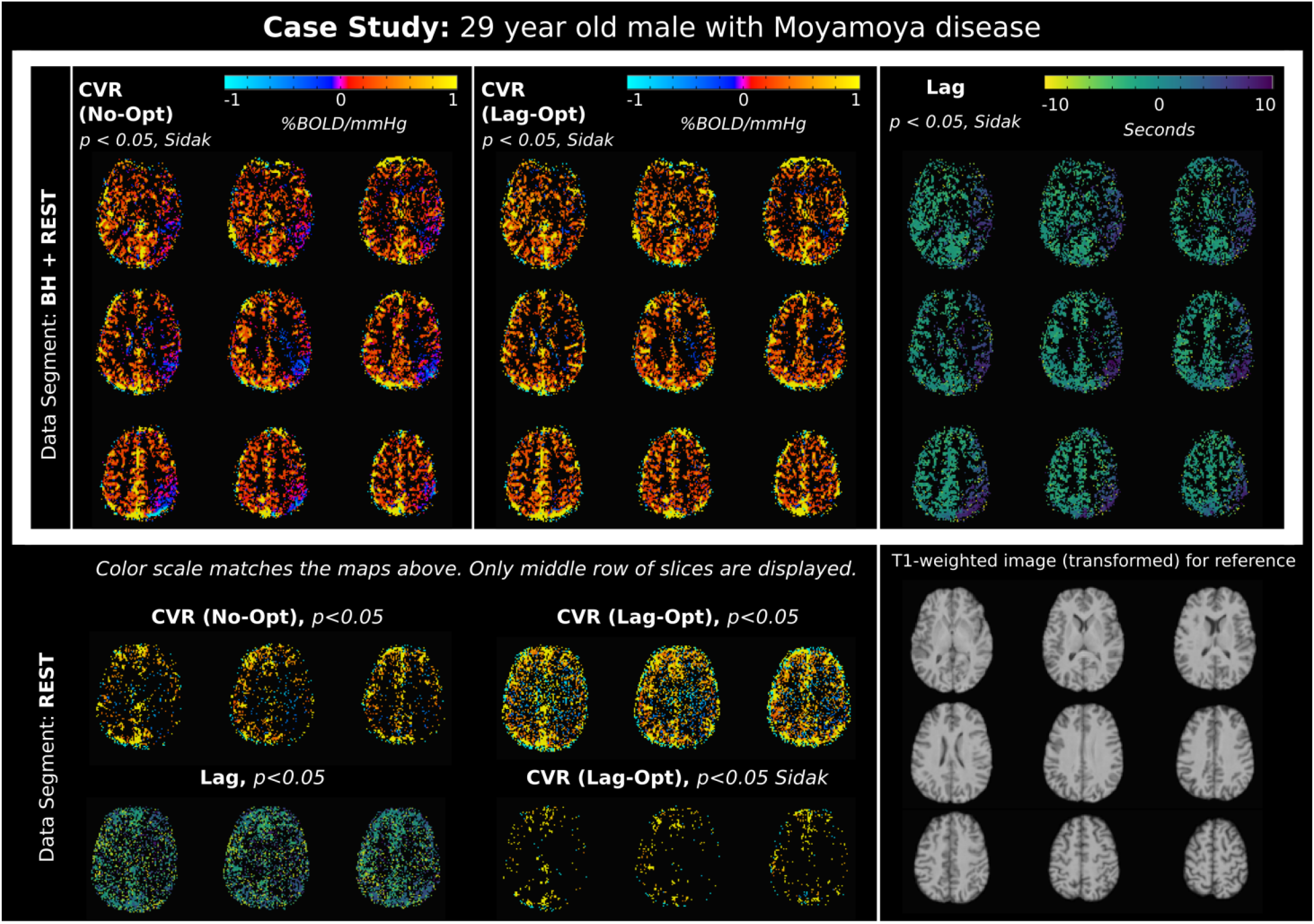
Maps of lag and CVR for a subject with Moyamoya disease. The white box shows results for the BH+REST data, demonstrating clear clinical sensitivity of this protocol. In order to visually compare maps across the same voxels, all maps (CVR No-Opt, CVR Lag-Opt and Lag) are thresholded at p<0.05, Sidak corrected, based on the T statistics of the lag optimized analysis. The REST only results are shown for comparison at p < 0.05 uncorrected as well as the p<0.05 Sidak corrected which displays very few significant voxels. Slices from the T1-weighted image transformed to fMRI space are shown for reference. CVR (Lag-Opt) maps, and all lag maps, do not include voxels with optimum lags found at the boundary. Lag maps are not relative to GM median.

In a concurrent pediatric pilot study, five individuals with typical development and hemiparetic cerebral palsy completed a modified version of the CDB+REST protocol. Figure 10 shows P_ET_CO_2_ traces and average BOLD-fMRI signal across GM. Three primary scenarios of task compliance were observed. In Scenario 1, participants achieved hypocapnia as expected, evidenced by two consecutive decreases in P_ET_CO_2_ and BOLD-fMRI signals. In Scenario 2, P_ET_CO_2_ values were unreliable for several breathing cycles (indicated by missing values in Figure 10), yet there is possible evidence of two mild hypocapnia cycles in the BOLD-fMRI signal. In Scenario 3, P_ET_CO_2_ values were more reliable but participants did not appear to complete the task, evidenced by a lack of hypocapnia-induced P_ET_CO_2_ and BOLD signal decreases. These trends appear to be age-related, with scenario 1 primarily occurring in older participants (ages 15-21 years) and scenarios 2 and 3 in the youngest participants (7-12 years).

**Figure 10.**
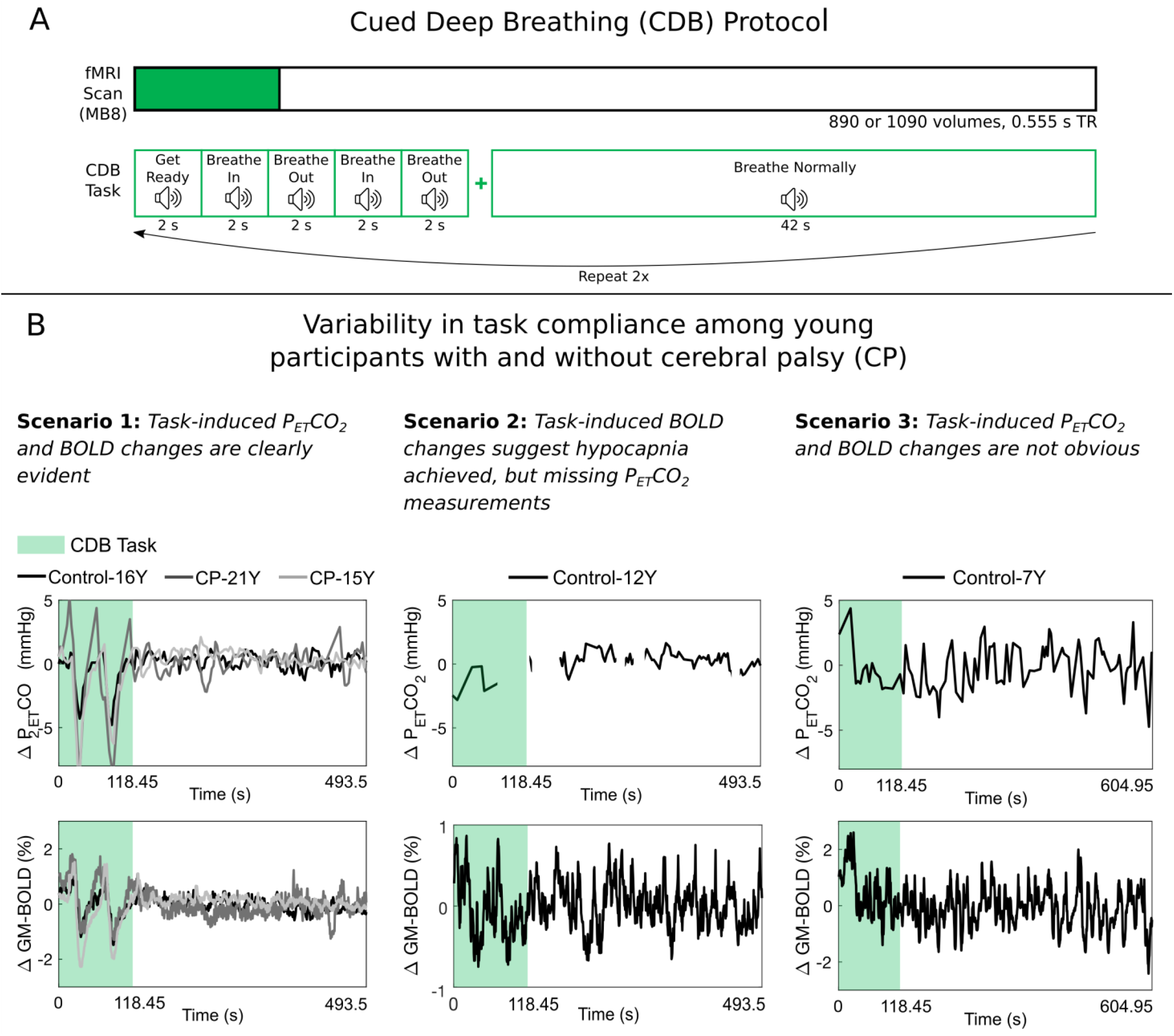
Task compliance for an adapted cued deep breathing (CDB) protocol used in a cohort of controls and individuals with hemiparetic cerebral palsy (CP). Panel (A) shows the protocol, which had the same task timings as the other CDB datasets in the manuscript, but with auditory cues, instead of visual. 20 s of rest were included at the start of the task, and 10 volumes (5.55 s) of data were removed to account for steady-state. Panel (B) shows unshifted P_ET_CO_2_ traces (mmHg change from baseline) and GM-BOLD traces (% change from mean) for 5 subjects. The missing values in the P_ET_CO_2_ traces in Scenario 2 indicate that the signal was unreliable at those times. Note the different axes limits for the GM-BOLD plot in Scenario 2. These subjects represent the primary age-related variations in task compliance observed during the CDB protocol.

## 4. DISCUSSION

Adding a simple, 2–3-minute breathing task to the beginning of a resting-state scan vastly improves our ability to model CO_2_ effects (both amplitude and timing) in BOLD-fMRI data. This modified scan protocol produces maps of physiological parameters that may contribute to our understanding of healthy and pathological cerebrovascular function, whilst maintaining an extended “resting state” period as required for studying intrinsic brain fluctuations and connectivity. This hybrid protocol has clear and direct applications for CVR mapping in both research and clinical settings, as well as wider applications for fMRI denoising and interpretation. Our analysis produces maps of both the amplitude (CVR) and timing (hemodynamic lag) of the BOLD fMRI response to CO_2_ by systematically shifting the P_ET_CO_2_ regressor to optimize the model fit. In any fMRI dataset, this optimization inherently increases GM CVR values and fit statistics. We have shown that the inclusion of breathing task data further increases the number of voxels in the brain that have a significant relationship between P_ET_CO_2_ and BOLD-fMRI signals, and improves our confidence in the plausibility of voxel-wise hemodynamic lag estimates.

A recent study [34] followed a very similar rationale to ours: to develop a practical non-gas inhalation method that is largely based in a resting-state scan but introduces breathing modulations to enhance fluctuations in P_ET_CO_2_. They compared CVR maps from resting-state scans, scans with intermittent breathing modulations throughout the scan period (for a duration of 12 seconds after 30-60 s of free breathing), a breath-holding scan, and a CO_2_ gas inhalation scan. They showed that intermittent breathing modulations (6 seconds per breath) had a comfort level similar to resting-state, and the resultant CVR maps had a sensitivity and accuracy similar to maps derived from gas inhalation methods. Hemodynamic lag was not a part of their assessment or comparisons. Their results are consistent with our conclusions that adding short periods of guided breathing is favorable over a resting-state scan for CVR mapping, and they have validated their design by comparing to a more gold standard CO2-inhalation approach. However, their breathing task and resting-state portion cannot be separated into two sections, precluding these data from being used for typical resting-state or functional connectivity analyses.

One reason that including breathing tasks benefits CVR mapping is simply that the induced variation in P_ET_CO_2_ amplitude is large compared to the smaller fluctuations during undirected free breathing (see Figure 2). Importantly, the amplitude of these natural fluctuations during resting-state will vary across subjects, and potentially across study populations [91]. If the BOLD fluctuations related to P_ET_CO_2_ changes are small in amplitude, it can be hard to distinguish them from other physiological, artefactual or neuronally-driven fluctuations that occur at (or aliased into) the same low-frequencies [45]–[49]. If these other fluctuations are of similar or greater magnitude to the low-frequency fluctuations induced by P_ET_CO_2_, this may result in an fMRI time-course poorly coupled to P_ET_CO_2_. A study using gas inhalation as a hypercapnic stimulus concluded that a change of at least 2 mmHg above a subject’s baseline P_ET_CO_2_ is necessary to evaluate hemodynamic impairment [92]. In our acquisitions, both BH and CDB tasks clearly produced changes above 2 mmHg, whereas this was not the case for all subjects during the REST segments (see Figure 2). Even if a subject performs the BH or CDB task only partially (i.e., shorter hold for the BH, shallower breaths for the CDB), they are likely to still surpass this 2 mmHg change. Furthermore, there is evidence that despite variability in BH performance, robust and repeatable CVR maps can still be obtained when modelling with P_ET_CO_2_ regressor [64], [93], which represents what the subject actually achieved and not simply the intended stimulus.

To accurately model CVR we must account for hemodynamic lags since measurement delays in gas sampling, arterial transit times to the brain’s vascular territories, and local vasodilatory dynamics all impact the temporal relationship between our model of the vasodilatory stimulus (P_ET_CO_2_) and the BOLD fMRI timeseries. Characterizing hemodynamic lag at the voxel level is both challenging and necessary for correct physiologic interpretation of the data. A previous study mapping CVR with resting-state data discussed how they were unable to obtain voxel-wise delays due to the resultant CVR maps being noisy, and they acknowledge that the regional CVR deficits they report in Moyamoya patients may reflect both reduced CVR and longer blood transits [32]. We report similar challenges in our data (Figure 3, Figure 6, Supp. Figure 1), showing more variable and less physiologically plausible lag distributions across GM in the data modelled with only resting-state segments. Furthermore, when simply performing a cross-correlation between the P_ET_CO_2_hrf regressor and GM BOLD-fMRI time-series, we observe several extreme lag values in resting-state data (Figure 4). The incorporation of the short breathing tasks results in more sensible lag distributions and cross-correlation results. Comparing our lag values to previous literature is challenging due to the multitude of experimental set-ups, analysis approaches, and ways of summarizing these types of data. Nevertheless, it is valid to compare normalized lag values (lag values relative to a tissue average) and compare variability in lag, where possible. Here, we focus on BH+REST and CDB+REST lag distributions, considering REST distributions are challenging to summarize (Figure 3). The range and variability of the lag values we report in GM show the majority of lag values (~68% based on one standard deviation, Supp. Table 1) are within 6 seconds of the GM median (Figure 6, Supplementary Figure 1); therefore, to capture the majority of GM lags a range of 12 seconds from the GM median may be appropriate. This broadly agrees with previous work using respiration derived or P_ET_CO_2_ regressors [43], [50], [57], [58], [61], [63], [64]. Many of these previous reports also see similar regional trends in relative lag values (Supplementary Figure 1): earliest responses in subcortical GM and later responses in cerebellar GM and posterior brain regions.

Despite much previous literature reporting a summary CVR value, our results clearly show that analysis choices in summarizing voxel-wise CVR values are not trivial. The distributions of GM CVR values with and without lag optimization, and with different levels of statistical thresholding (Figure 7, Sup. Figures 2-7), illustrate that it is not strictly valid to extract a central tendency value from a distribution of positive and negative CVR values together, after lag optimization or any statistical thresholding. We therefore chose to summarize positive and negative CVR values separately, not least because true negative CVR responses represent very different physiological mechanisms [86], [87]. We expect positive CVR values in GM, and it is noteworthy that the three resting-state segments show a greater number of negative CVR values compared to the segments that include BH or CDB tasks, possibly suggesting a greater relative contribution of noise sources and a less successful CVR and lag estimation. The final summative CVR value across a tissue type will also depend on whether one has applied lag optimization and/or applied statistical thresholding. Therefore, we chose to display CVR values resulting from multiple analysis options in Figure 8 and Supplementary Figure 7. With no statistical thresholding, the positive GM CVR amplitudes for No-Opt and Lag-Opt (Figure 8 and Supplementary Table 2) agree with the range seen in previous literature modelling CVR with breathing tasks or resting-state, when expressed in units of %BOLD/mmHg [34], [42], [54], [58], [63], [93], [94]. Unexpectedly, the REST segments showed higher GM CVR values than the task segments after lag optimization and following the strictest thresholding, particularly for REST_CDB_ (Figure 8, Supplementary Figure 7). However, a higher CVR value will not always suggest a more accurate or more representative tissue estimate. Low frequency fluctuations other than P_ET_CO_2_, motion artifacts and large vessel signals may influence the lagged fitting procedure, as discussed. There were generally more negative CVR values in REST segments, so positive GM CVR estimates are also summarized over a smaller number of voxels that do not fully describe GM tissue. It is also important to note that the Sidák correction is likely too strict of a correction for multiple comparisons as it assumes independence, which is not the case when running the GLM multiples times with slightly shifted P_ET_CO_2_ regressors.

### 4.1. BH vs. CDB

This study was not designed for an effective comparison between BH or CDB, yet some trends can be noted upon. Adding either a short BH or CDB task to a resting segment brings clear advantages for CVR mapping, with similar CVR and lag results, however the BH addition did lead to plausible lag distributions and significant CVR effects more consistently across subjects. Direct comparison of these two tasks is potentially biased due to the BH task being slightly longer, including 3 cycles of hypercapnia versus 2 cycles of hypocapnia for the CDB task. However, it is difficult to match both the length of the breathing task, and the number of cycles: the ways in which we achieve hypocapnia (increasing the rate and depth of breathing) and hypercapnia (apnea) are very different, and the timing dynamics of the resulting blood gas changes are different [95]–[97]. A small amount of previous evidence shows that CDB and BH derived CVR responses generally have comparable amplitudes and timings [50], however possibly not in all brain regions [94] and they can be modulated differently by baseline vasodilation [94]. Furthermore, these tasks induce different motion confounds [50] and potentially different subject compliance demands. In general, our results suggest they may be used interchangeably in healthy controls. However, a more thorough comparison of the pros and cons of these breathing tasks, in healthy and clinical cohorts, is a worthwhile focus for future research. Importantly, vasoconstrictive and vasodilatory responses can be differentially affected in pathology [98], so different tasks may be appropriate for different clinical populations and study goals.

It is important to discuss the general limitations of using breathing tasks to model CVR, compared to resting-state. They do require more compliance from subjects, introduce neural activity due to the need for visual or auditory stimuli, and often exhibit motion effects correlated with task timings [50]–[52], [64]. We performed volume realignment and included the resultant motion parameters within the GLM, but there is evidence this is not sufficient to remove motion effects [52], [64], [99]. Further research should look into optimizing breathing task designs to minimize task induced motion artifacts whilst still maintaining P_ET_CO_2_ changes of sufficient amplitude. Collecting multi-echo BOLD fMRI data, and applying spatial ICA-denoising analyses, is one widely adopted strategy that can be applied to remove motion artifacts and has been applied to the modelling of CVR effects in BH paradigms [56], [64]. A final consideration is the choice of HRF in breathing task data; here we used a canonical HRF to convolve with all the P_ET_CO_2_ time-series, before creating the multiple shifted P_ET_CO_2_ regressors. There is evidence that the BOLD signal may have a different shape and timing response depending on whether the P_ET_CO_2_ is resting fluctuations, or within a hypocapnic or hypercapnic range [57], and that physiological responses functions can vary across subjects, brain regions and sessions [100], [101]. Therefore, one response function might not be fully optimum in capturing all the P_ET_CO_2_ induced BOLD changes within our hybrid design, yet there weren’t sufficient repeats of the BH or CDB cycles to characterize an impulse response function from these data.

### 4.2. Clinical applications

We chose to analyze and present our results in single subject space, partially to demonstrate the potential utility of this method for individual subjects, such as clinical cases, or individual fMRI denoising. We report an incidental finding of Moyamoya disease, which was based on our inspection of the hemodynamic lag maps constructed from our lagged-GLM approach. This pathology was only visually obvious when modeling lag and CVR with breathing task data. Regional deficits in CVR are used to guide surgical and treatment decisions in cases of Moyamoya [102], with normalization of CVR often seen after surgery. Delayed blood transits are also widely reported in cases of Moyamoya [61], [89], [103] which is expected due to the narrowing of blood vessels. Consistent with our results, negative CVR responses are often observed in cases of Moyamoya [88]–[90] and commonly interpreted as the vascular steal phenomenon. In the case of vascular steal, negative CVR is the result of a redistribution of blood flow from regions without remaining cerebrovascular reserve to regions with preserved vasodilatory capacity. Previous work with cases of Moyamoya have shown a clear relationship between blood arrival times and CVR amplitudes, with the longest arrival times having the lowest and most negative CVR (e.g., [89], [103]. From our data, it is possible to conclude there is no evidence of vascular steal, considering the normalization of CVR after lag-optimization, however other work suggests that negative CVR is likely a combination of a steal phenomenon and delayed local (positive) reactivity [89]. We would need to apply our technique more systematically across a larger sample to better characterize these subtleties of Moyamoya pathology, yet what our case study clearly demonstrates is the importance of considering both amplitude and timing when modelling CVR function in pathology. CVR deficits in Moyamoya are most commonly investigated with gas inhalation or contrast; although there is a much smaller collection of literature using breath-hold or resting-state, the interest in these non-invasive and practical approaches is growing [35], [61], [90], [104], [105].

Example data from a separate pediatric pilot study were included to evaluate the feasibility of our hybrid protocol in clinical cohorts with task compliance challenges (e.g., children). Breathing tasks are an attractive alternative to more invasive vasoactive stimuli; BH tasks have been used successfully in typically developing children [106] and in those with Moyamoya [90]. We report the first instance of a CDB task in a pediatric cohort and observed variable success in the quality of P_ET_CO_2_ recordings and achieved hypocapnia. These results indicate that further optimization of the breathing task portion of the hybrid design may be necessary in less-compliant cohorts. Low quality P_ET_CO_2_ traces may be caused by periods of mouth breathing and could be addressed by using a mask rather than a nasal cannula. It may be necessary to adapt breathing task instructions for clarity and utilize practice sessions, tailored to the population, to ensure participants understand and comply with the task before entering the scanner. There are clear next steps to address the feasibility limitations observed in our hybrid design and, once these are met, this method has promise as a practical yet robust tool to study typical and atypical neurovascular development.

### 4.3. Applications for fMRI denoising

Based on the mechanism of neurovascular coupling, the BOLD contrast is widely used as a surrogate measure for neural activity [107]. However, there are many non-neural factors that can affect the BOLD signal, with multiple strategies and algorithms to mitigate and remove these sources of noise, [31], [45], [47], [108]–[112]. We considered how resting-state fMRI scans are commonly deployed in neuroimaging research, and designed our protocol accordingly. Although the main focus of this work is practical CVR mapping, CO_2_ fluctuations have been shown to be a physiological confound in resting-state fMRI studies of neural activity, and it is challenging to meaningfully separate out neural connectivity and vascular connectivity with fMRI data [51], [111], [113]–[116]. Removing CO_2_ effects from fMRI via nuisance regression is not yet routine practice, in part because it is challenging to robustly characterize these relationships in resting-state fMRI. By appending a short breathing task to the start of a resting-state scan, the resting-state portion could still be analyzed separately, gaining more outputs from one dataset, with improved denoising from robust modelling of CO_2_ effects.

However, there is a possibility that the breathing task may somehow modulate the resting-state portion of the scan similar to after effects seen after other sensory and cognitive tasks [117], [118]. We performed exploratory analyses to compare the three REST segments and assess any systematic differences caused by preceding breathing tasks. We created maps of amplitude of low-frequency fluctuation (ALFF), fractional ALFF (fALFF), and local functional connectivity density (LFCD) for each subject and each REST segment using AFNI 3dRSFC and 3dLFCD functions, respectively. Spatial correlations were performed between pairs of single-subject maps. For ALFF, group means ± stdev were: 0.92±0.04 (REST, REST_BH_), 0.93±0.04 (REST, REST_CDB_) and 0.93±0.03 (REST_BH_, REST_CDB_). For fALFF: 0.87±0.08 (rest, REST_BH_), 0.91±0.03 (REST, REST_CDB_), 0.90±0.05 (REST_BH_, REST_CDB_). For LFCD: 0.62±0.21 (REST, REST_BH_), 0.58±0.25 (REST, REST_CDB_), 0.66±0.19 (REST_BH_, RESTCDB). We also performed a voxel-wise ANOVA analysis (3dANOVA, AFNI) to test if there was a significant difference between the three rest segments for each type of map. Setting a threshold of p<0.05 (FDR corrected), we observed no significant voxels for ALFF, fALFF or LFCD (main effect and pairwise comparisons). These results show no clear systematic differences between the three REST segments for ALFF, fALFF or LFCD. However, these analyses cannot provide direct evidence for the null hypothesis, and future work with study designs tailored towards this question, and with bigger samples, should investigate this further.

### 4.4. Recommendations and practical considerations

In this paper, some analysis choices were driven by the comparison between breathing task data segments and resting-state only data segments. If a researcher was to use this BH+REST or CDB+REST design as shown in Figure 1, we recommend:

- Collect continuous CO_2_ recordings before and after your scan window, up to the maximum shift you want to consider in your modeling, to avoid interpolation or trimming of data after shifting.
- If deciding between a BH or CDB task, we’d recommend BH due to more extensive literature using this task for CVR mapping [14], [15].
- Model CVR using the entire dataset. We cut the TASK+REST segment to 8 minutes in order to make a fair comparison with the 8–minute REST segments. Therefore, our CVR and lag maps are from modelling with ~2.5 minutes of task data and 5.5 minutes of resting; map quality and fit statistics would likely improve with more data.
- If there is a large offset between the P_ET_CO_2_ trace and the GM fMRI signal, perhaps due to a measurement delay (e.g., long sample line from scanner to control room), first perform a bulk-shift (cross-correlation between a mean GM fMRI time-series and P_ET_CO_2_ time-series) before voxel-wise optimization [63], supported by Figure 4.
- Use the smallest range of shifted P_ET_CO_2_ regressors as is appropriate for the physiology of your cohort. There are computational and statistical consequences to extending the shift range more than is necessary.
- When obtaining a summary CVR or lag metric for a tissue class, check the distributions to see if this is valid and appropriate.

### 4.5. Conclusions

This hybrid design may not the best approach for all CVR mapping experiments; if that is your only research focus, longer breathing tasks or gas inhalation methods are recommended. However, our proposed protocol involves a small addition to a typical resting-state scan, aiming to be practical and clinically feasible in situations where more invasive or complicated methods are not desirable. We have demonstrated that adding a short breathing task to the start of a resting-state fMRI scan improves the ability to model both the timing and amplitude of the CVR response, both crucially important for the accurate characterization of cerebrovascular function.

## Abbreviations

No-Opt: (No Optimization)
Lag-Opt: (Lag Optimization)
HRF/hrf: (Hemodynamic Response Function)
PaCO_2_: (Partial pressure arterial CO_2_)
P_ET_CO_2_: (Partial pressure of End Tidal CO_2_)
BH: (Breath Holding)
CDB: (Cued Deep Breathing)
GLM: (Generalized Linear Model)
SBRef: (Single Band Reference image)
GM: (Grey matter)
Cerebrovascular reactivity: (CVR)

## 5. ACKNOWLEDGMENTS

This research was supported by the Eunice Kennedy Shriver National Institute of Child Health and Human Development of the National Institutes of Health under award number K12HD073945. The pediatric dataset and cerebral palsy dataset were collected with support of National Institutes of Health award R03 HD094615-01A1. The authors would like to acknowledge Marie Wasielewski and Carson Ingo for their support in acquiring these data.

K.Z. was supported by an NIH-funded training program (T32EB025766). S.M. was supported by the European Union’s Horizon 2020 research and innovation program (Marie Skłodowska-Curie grant agreement No. 713673), a fellowship from La Caixa Foundation (ID 100010434, fellowship code LCF/BQ/IN17/11620063) and C.C.G was supported by the Spanish Ministry of Economy and Competitiveness (Ramon y Cajal Fellowship, RYC-2017-21845), the Basque Government (BERC 2018-2021 and PIBA_2019_104) and the Spanish Ministry of Science, Innovation and Universities (MICINN; PID2019-105520GB-100).

## CrediT statement

Rachael Stickland: Conceptualization, Methodology, Software, Formal Analysis, Investigation, Data Curation, Writing (OD), Writing (RE), Visualization, Project Administration. Kristina Zvolanek: Conceptualization, Software, Investigation, Formal Analysis, Writing (RE), Visualization. Stefano Moia: Methodology, Writing (RE).

Apoorva Ayyagari: Investigation. César Caballero-Gaudes: Methodology, Writing (RE). Molly Bright: Conceptualization, Methodology, Software, Investigation, Resources, Writing (RE), Supervision, Project Administration, Funding Acquisition.

The authors would also like to thank Kevin Murphy for the basis of the code that creates the physiological regressors, and personnel at the Center for Translation Imaging (Northwestern Radiology) for support with study set-up.

## 6. COMPETING FINANCIAL INTERESTS

The authors declare no competing financial interests.

## Notes

### Competing Interest Statement

The authors have declared no competing interest.

### Summary of Updates

Small wording corrections.

