## Supplementary Data for "A practical modification to a resting state fMRI protocol for improved characterization of cerebrovascular function"

### Supplementary Material

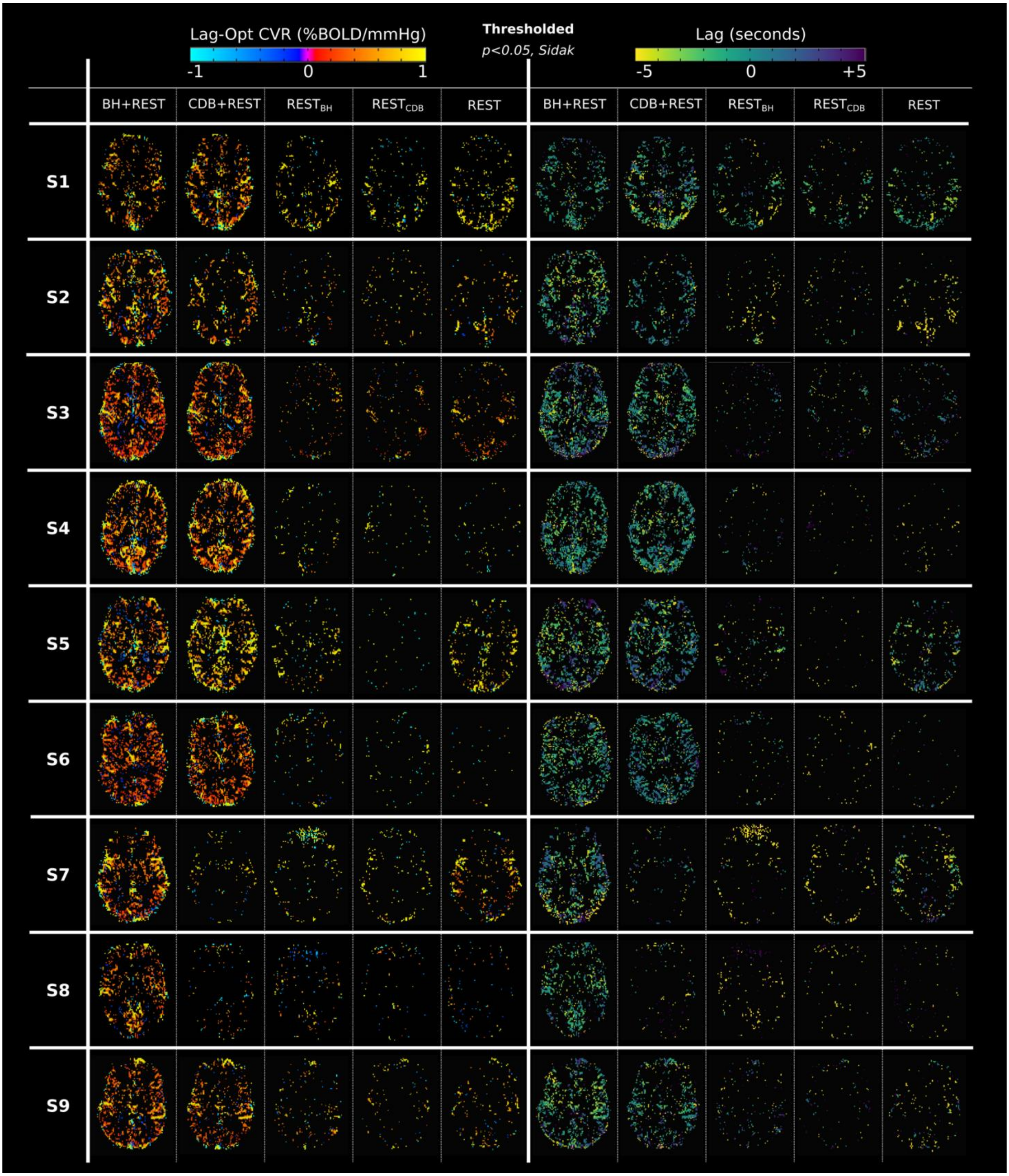


**Supplementary Figure 1.** Maps of lag optimized CVR (left columns) and of lag (right columns) displayed for each data segment and each subject (S1-S9). All maps are shown with the same statistical threshold, p<0.05 with Sidak correction. All maps do not include voxels with optimum lags found at the boundary.

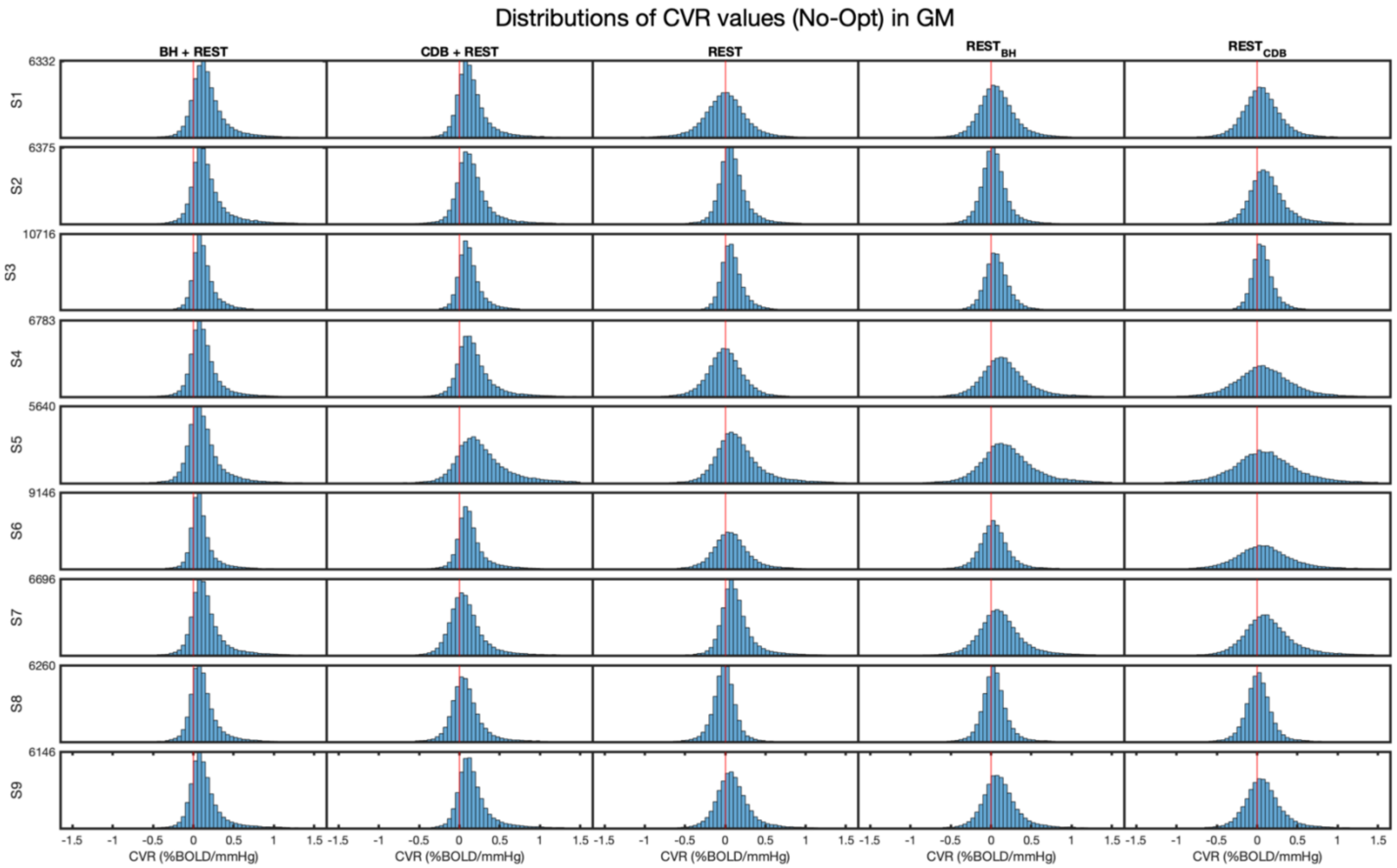


**Supplementary Figure 2.** Distributions of No-Opt CVR values in GM, for each subject (S1-S9). No statistical thresholding has been applied therefore all GM voxels are included. The y-axes show the histogram frequency count. The axes are scaled to the same maximum for each row, and these maximums are shown in the first column only. The red lines indicate the position of ’0’ on the x-axes which shows the CVR bins.

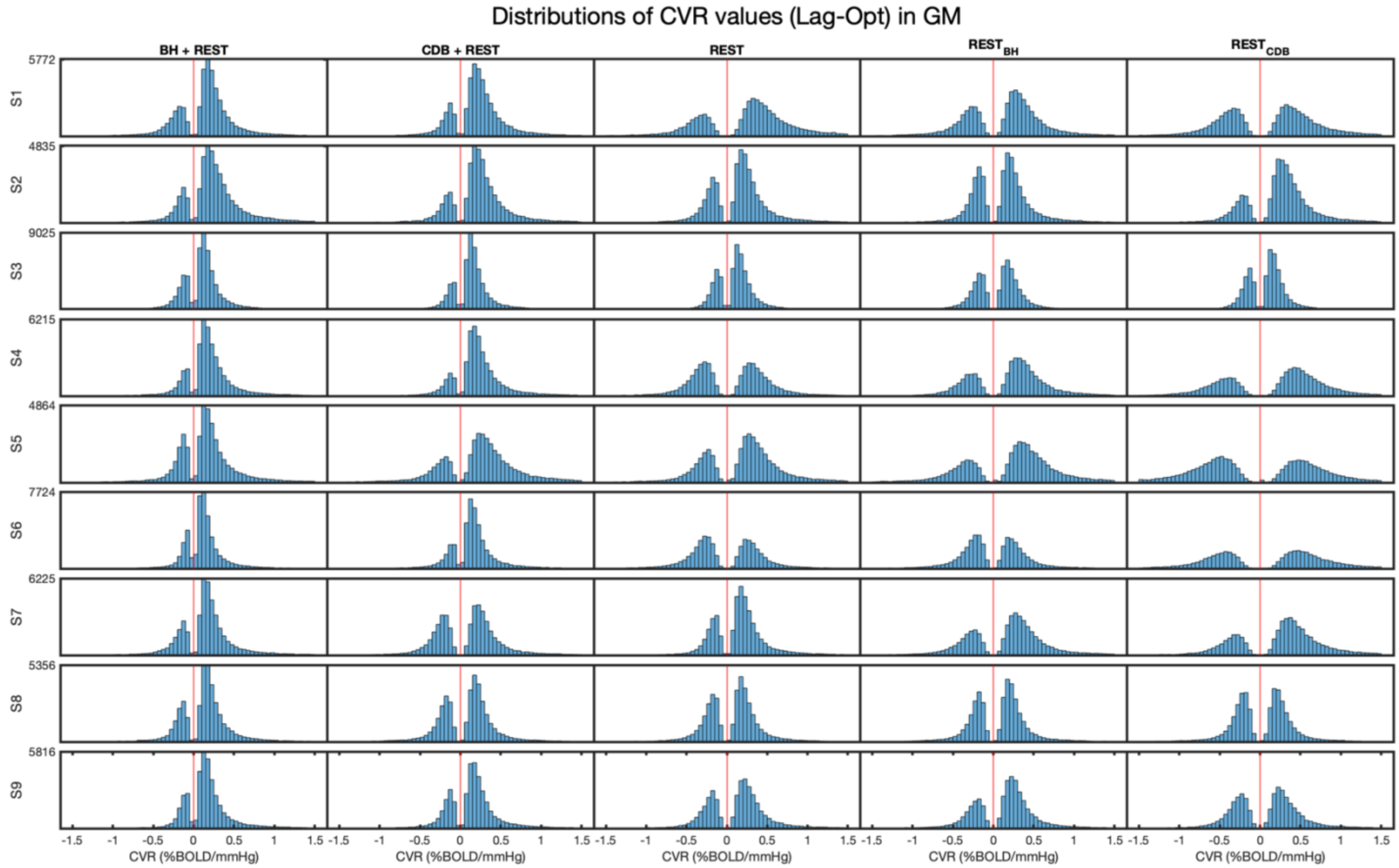


**Supplementary Figure 3.** Distributions of Lag-Opt CVR values in GM, for each subject (S1-S9). No statistical thresholding has been applied therefore all GM voxels are included. The y-axes show the histogram frequency count. The axes are scaled to the same maximum for each row, and these maximums are shown in the first column only. The red lines indicate the position of ’0’ on the x-axes which shows the CVR bins.

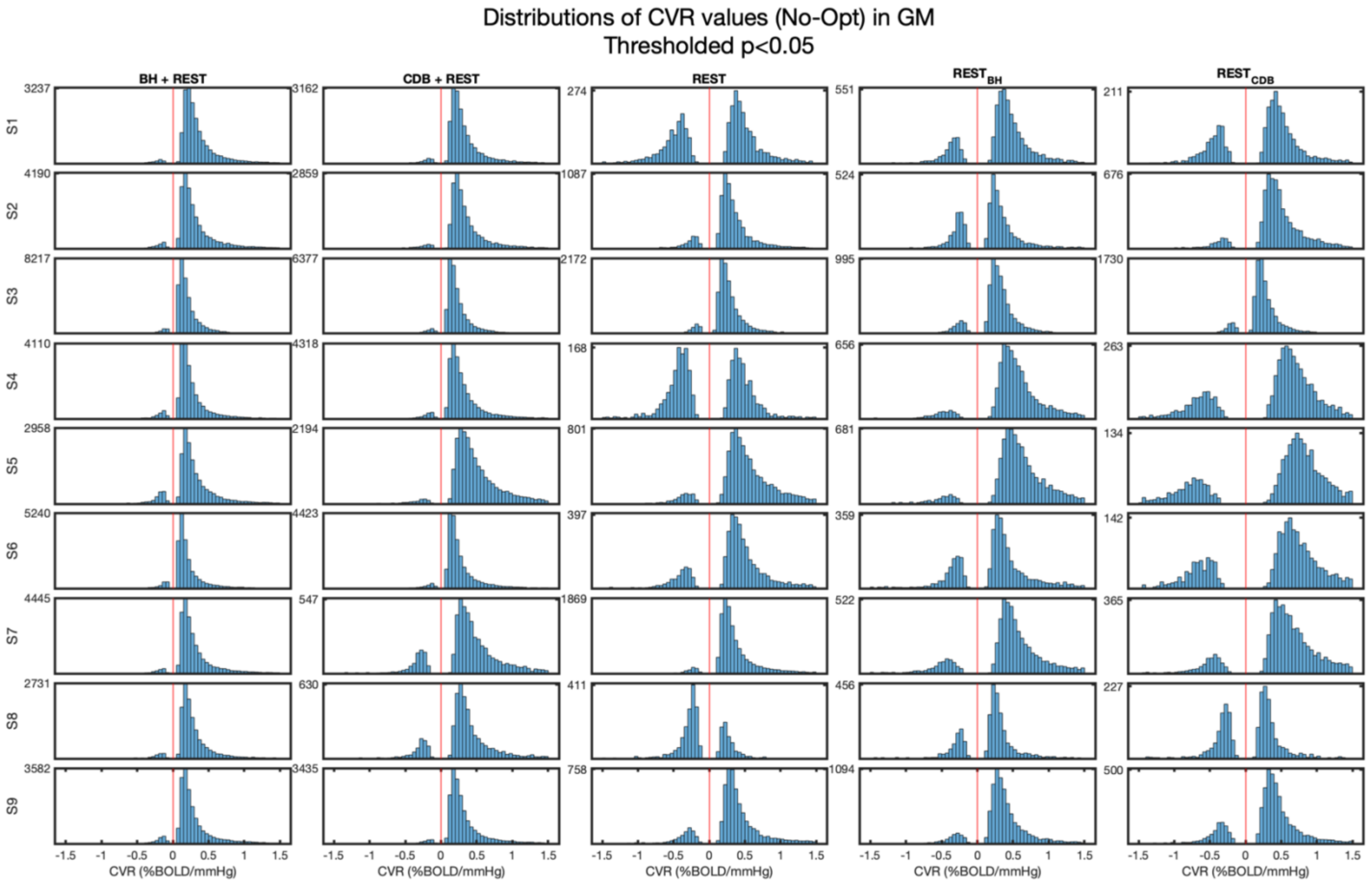


**Supplementary Figure 4.** Distributions of No-Opt CVR values in GM, for each subject (S1-S9). Only voxels that were significant (at $p<0.05$) were included. Therefore, different numbers of GM voxels remain for each data segment and each subject. The y-axes show the histogram frequency count. The number shown at the top left-hand corner of each plot is the maximum frequency count for that plot. The red lines indicate the position of ’0’ on the x-axes which shows the CVR bins.

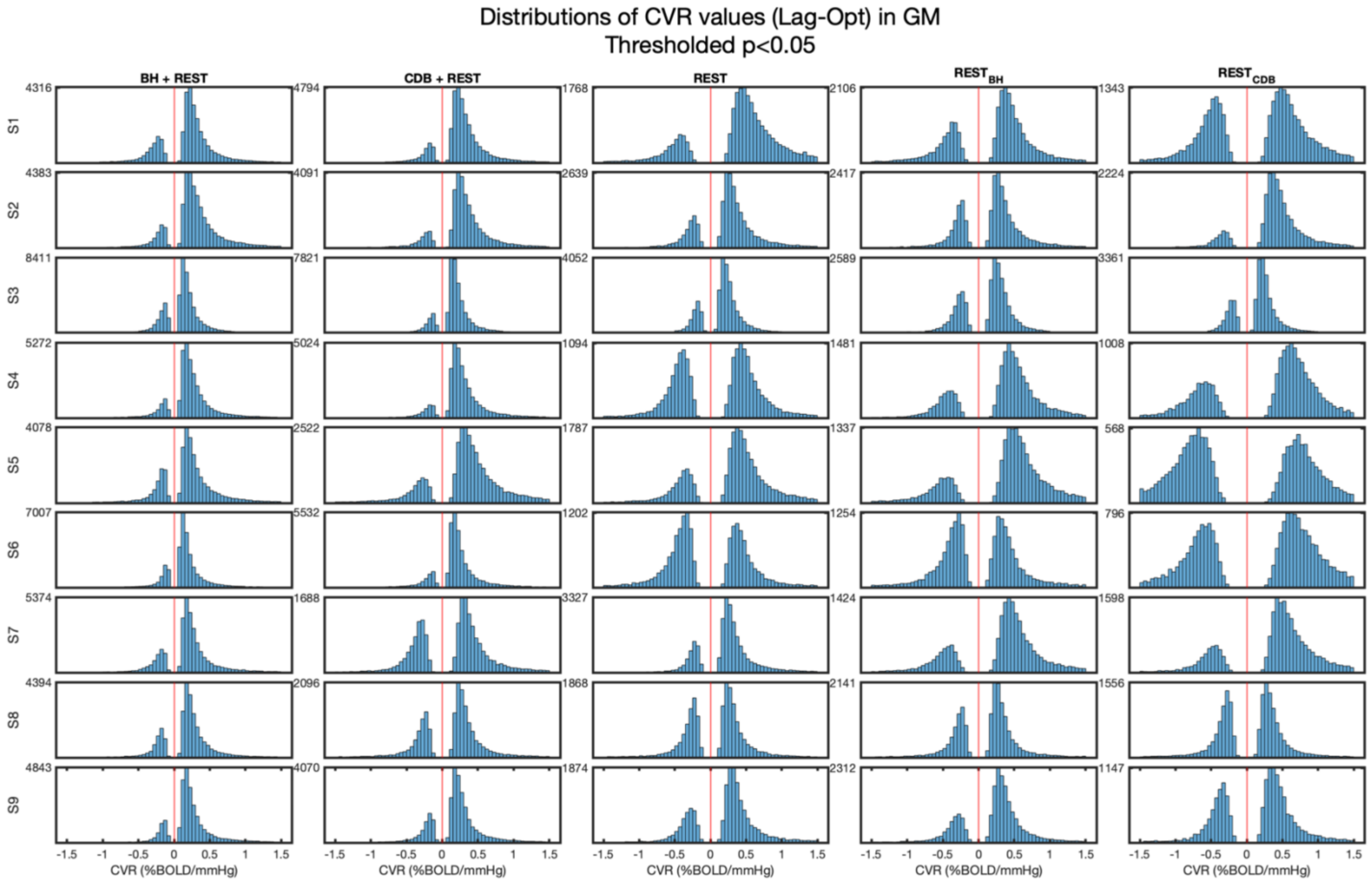


**Supplementary Figure 5.** Distributions of Lag-Opt CVR values in GM, for each subject (S1-S9). Only voxels that were significant (at p<0.05) were included. Therefore, different numbers of GM voxels remain for each data segment and each subject. The y-axes show the histogram frequency count. The number shown at the top left-hand corner of each plot is the maximum frequency count for that plot. The red lines indicate the position of ’0’ on the x-axes which shows the CVR bins.

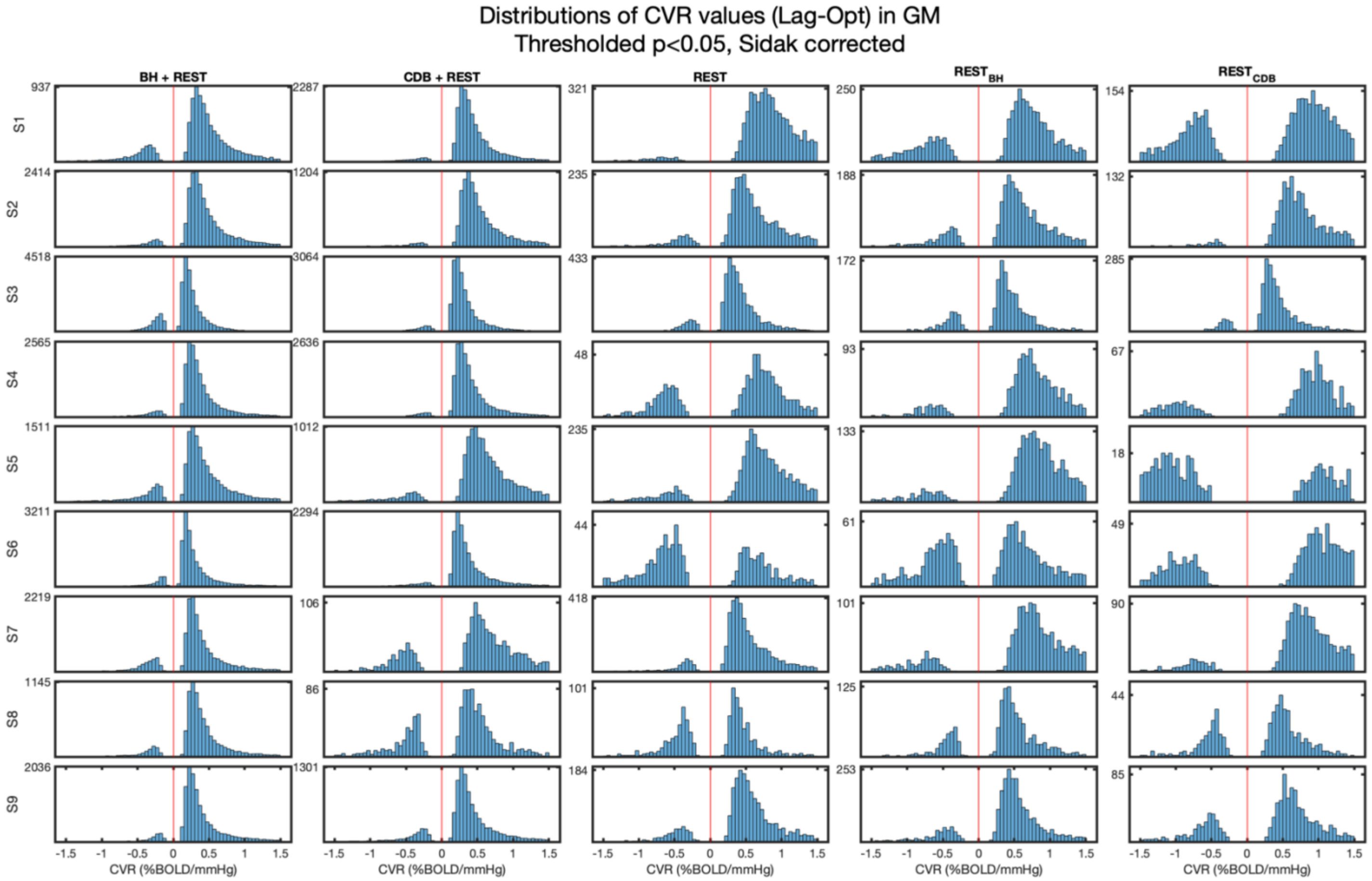


**Supplementary Figure 6.** Distributions of Lag-Opt CVR values in GM, for each subject (S1-S9). Only voxels that were significant (at $p<0.05$, Sidak corrected) were included. Therefore, different numbers of GM voxels remain for each data segment and each subject. The y-axes show the histogram frequency count. The number shown at the top left-hand corner of each plot is the maximum frequency count for that plot. The red lines indicate the position of ’0’ on the x-axes which shows the CVR bins.

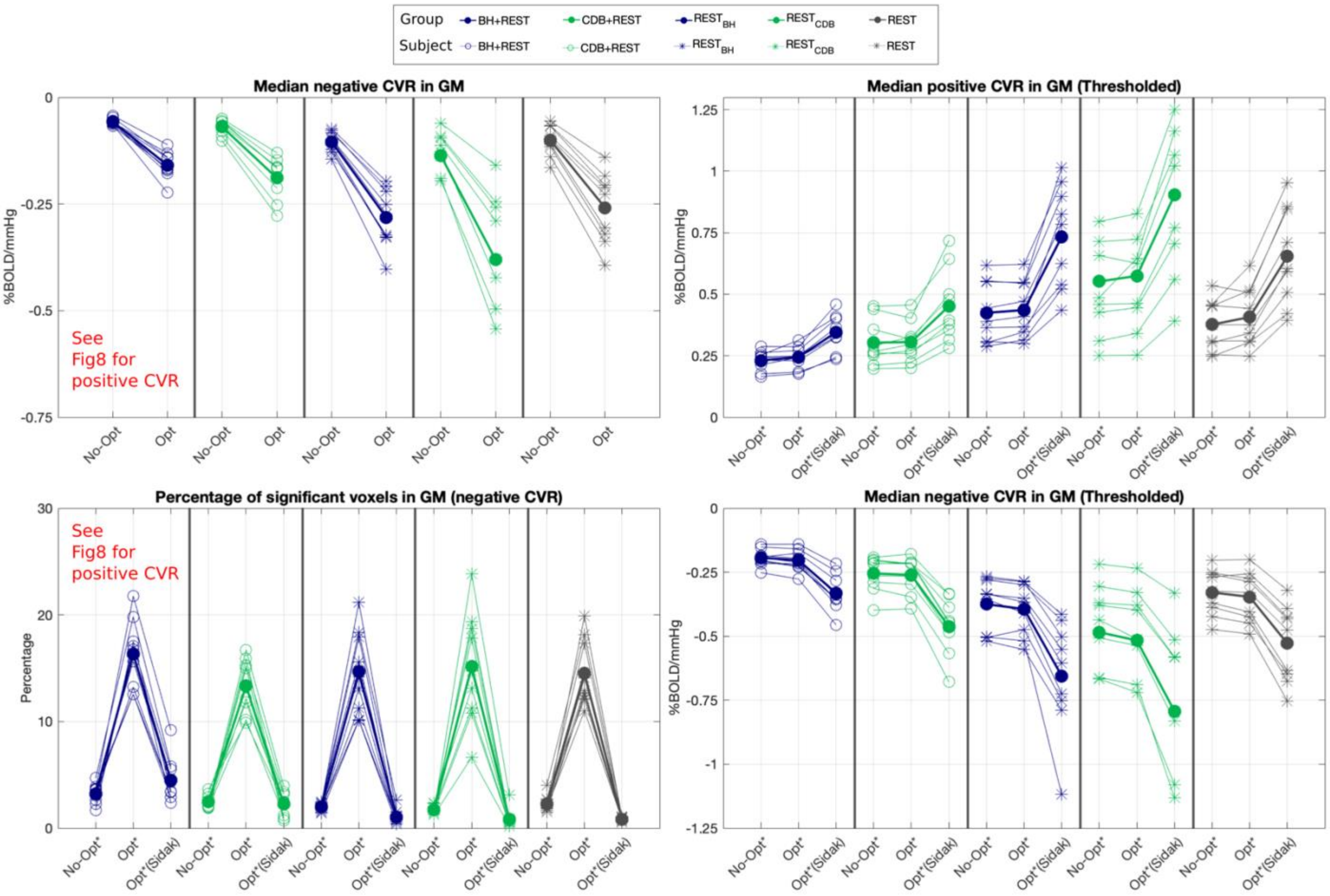


**Supplementary Figure 7.** Comparing GM summary metrics across the five data segments. The legend corresponds to the column structure in each of the four panels; both single subject data and group means are shown. *Top Left:* Median negative CVR across all GM voxels for non-optimized (No-Opt) and lag optimized (Lag-Opt) analyses. *Bottom Left:* Percentage of significant negative fits in GM for different analyses and levels of thresholding. *Right panels*: Median positive CVR (*top*) and negative CVR (*bottom*) in GM for No-Opt and Lag-Opt for different thresholding levels: [ * ] indicates thresholding at $p<0.05$ and [ (*Sidak) ] indicates thresholding at $p<0.05$ with Šidák correction. Median CVR values that were taken from statistically thresholded distributions also did not include voxels where lag was found at the boundary condition.

| Lag values | All GM | | | Regional (relative to all GM median) | | | Earliest to latest response |
|---|---|---|---|---|---|---|---|
| | Median | Mean | Stdev | Cerebellar GM median | Cortical GM median | Subcortical GM median | |
| **BH+REST** | | | | | | | |
| S1 | -1.8 | -1.91 | 5.92 | 2.1 | 0 | 0.3 | *Cortical, subcortical, cerebellar* |
| S2 | -4.8 | -3.60 | 5.79 | 2.1 | -0.6 | -1.2 | *Subcortical, cortical, cerebellar* |
| S3 | -0.6 | -0.75 | 6.07 | 2.1 | 0.3 | 0 | *Subcortical, cortical, cerebellar* |
| S4 | -3 | -2.49 | 5.49 | 1.2 | 0 | 0 | *Subcortical & cortical, cerebellar* |
| S5 | -0.9 | -0.79 | 6.49 | 3.75 | 0 | -1.5 | *Subcortical, cortical, cerebellar* |
| S6 | -3 | -2.28 | 6.07 | 1.5 | 0 | -0.6 | *Subcortical, cortical, cerebellar* |
| S7 | -1.2 | -1.60 | 6.35 | 1.5 | 0.3 | 0.15 | *Subcortical, cortical, cerebellar* |
| S8 | -2.7 | -1.47 | 6.87 | 2.4 | -0.3 | -0.3 | *Subcortical & cortical, cerebellar* |
| S9 | -1.8 | -1.59 | 5.81 | 1.5 | 0.3 | -0.3 | *Subcortical, cortical, cerebellar* |
| Mean | -2.20 | -1.83 | 6.10 | 2.02 | 0.00 | -0.38 | ***Most common: Subcortical, cortical, cerebellar*** |
| Stdev | 1.32 | 0.89 | 0.42 | 0.76 | 0.30 | 0.61 | |
| **CDB+REST** | | | | | | | |
| S1 | -4.8 | -4.14 | 5.70 | 1.8 | -0.3 | -0.9 | *Subcortical, cortical, cerebellar* |
| S2 | -3.9 | -2.77 | 6.24 | 1.5 | 0 | -0.3 | *Subcortical, cortical, cerebellar* |
| S3 | 0.3 | -0.37 | 6.04 | 2.1 | 0 | -1.2 | *Subcortical, cortical, cerebellar* |
| S4 | -2.1 | -1.92 | 5.31 | 0.9 | 0 | -0.15 | *Subcortical, cortical, cerebellar* |
| S5 | -0.9 | -1.41 | 5.98 | 1.8 | 0 | -0.9 | *Subcortical, cortical, cerebellar* |
| S6 | -2.4 | -2.27 | 5.45 | 0.6 | 0.3 | -0.6 | *Subcortical, cortical, cerebellar* |
| S7 | -2.4 | -1.30 | 7.91 | 0.3 | 0 | 1.05 | *Cortical, cerebellar, subcortical* |
| S8 | -0.3 | -0.25 | 7.86 | 2.1 | 0.3 | -0.75 | *Subcortical, cortical, cerebellar* |
| S9 | -1.5 | -1.53 | 6.14 | 1.8 | 0.3 | 0.6 | *Cortical, subcortical, cerebellar* |
| Mean | -2.00 | -1.77 | 6.29 | 1.43 | 0.07 | -0.35 | ***Most common: Subcortical, cortical, cerebellar*** |
| Stdev | 1.64 | 1.20 | 0.95 | 0.67 | 0.20 | 0.75 | |

**Supplementary Table 1.** Lag values across different GM regions, shown for BH+REST and CDB+REST data segments, for each of the 9 subjects. Before medians were calculated across different parts of GM (cerebellar, cortical, subcortical) the values were changed to be relative to global GM median. Mean and standard deviation (Stdev) for all GM is also shown for reference. Voxels with lag values found at the boundary condition were not included in the calculations.

| Parameter across GM voxels | Direction | BH+REST | | CDB+REST | | REST | | $REST_{BH}$ | | $REST_{CDB}$ | |
|---|---|---|---|---|---|---|---|---|---|---|---|
| | | *Mean* | *Stdev* | *Mean* | *Stdev* | *Mean* | *Stdev* | *Mean* | *Stdev* | *Mean* | *Stdev* |
| Median CVR (No-Opt) | Positive | 0.14 | 0.02 | 0.17 | 0.05 | 0.15 | 0.04 | 0.17 | 0.06 | 0.20 | 0.07 |
| Median CVR (Lag-Opt) | Positive | 0.21 | 0.04 | 0.25 | 0.06 | 0.31 | 0.10 | 0.32 | 0.09 | 0.43 | 0.16 |
| Median CVR (No-Opt) p<0.05 | Positive | 0.23 | 0.04 | 0.30 | 0.09 | 0.38 | 0.10 | 0.42 | 0.12 | 0.55 | 0.22 |
| Median CVR (Lag-Opt) p<0.05 | Positive | 0.24 | 0.04 | 0.31 | 0.08 | 0.41 | 0.12 | 0.44 | 0.12 | 0.57 | 0.21 |
| Median CVR (Lag-Opt) p<0.05, Šidák | Positive | 0.35 | 0.07 | 0.45 | 0.15 | 0.65 | 0.20 | 0.73 | 0.21 | 0.90 | 0.31 |
| Median T-statistic (No-Opt) | Positive | 2.21 | 0.36 | 1.95 | 0.65 | 1.00 | 0.28 | 1.01 | 0.17 | 0.88 | 0.16 |
| Median T-statistic (Lag-Opt) | Positive | 3.19 | 0.38 | 2.82 | 0.59 | 2.03 | 0.19 | 1.99 | 0.13 | 1.93 | 0.15 |
| Percentage of significant fits (No-Opt) p<0.05 | Positive | 44.86 | 7.84 | 37.94 | 16.87 | 11.79 | 8.20 | 11.08 | 5.25 | 7.67 | 4.44 |
| Percentage of significant fits (Lag-Opt) p<0.05 | Positive | 60.23 | 5.31 | 53.64 | 15.50 | 33.45 | 9.40 | 31.90 | 5.78 | 28.98 | 7.57 |
| Percentage of significant fits (Lag-Opt) p<0.05, Šidák | Positive | 31.20 | 8.65 | 24.23 | 13.84 | 5.42 | 3.77 | 4.06 | 1.74 | 2.69 | 1.85 |
| Median CVR (No-Opt) | Negative | -0.06 | 0.01 | -0.07 | 0.02 | -0.10 | 0.04 | -0.10 | 0.02 | -0.14 | 0.05 |
| Median CVR (Lag-Opt) | Negative | -0.16 | 0.03 | -0.19 | 0.05 | -0.26 | 0.08 | -0.28 | 0.07 | -0.38 | 0.16 |
| Median CVR (No-Opt) p<0.05 | Negative | -0.19 | 0.03 | -0.26 | 0.07 | -0.33 | 0.09 | -0.37 | 0.11 | -0.48 | 0.19 |
| Median CVR (Lag-Opt) p<0.05 | Negative | -0.20 | 0.04 | -0.26 | 0.07 | -0.35 | 0.10 | -0.39 | 0.10 | -0.52 | 0.20 |
| Median CVR (Lag-Opt) p<0.05, Šidák | Negative | -0.33 | 0.07 | -0.46 | 0.11 | -0.53 | 0.15 | -0.66 | 0.22 | -0.79 | 0.32 |
| Median T-statistic (No-Opt) | Negative | -0.85 | 0.11 | -0.75 | 0.10 | -0.65 | 0.05 | -0.64 | 0.04 | -0.61 | 0.03 |
| Median T-statistic (Lag-Opt) | Negative | -2.32 | 0.21 | -2.01 | 0.16 | -1.77 | 0.06 | -1.76 | 0.13 | -1.76 | 0.14 |
| Percentage of significant fits (No-Opt) p<0.05 | Negative | 3.24 | 0.92 | 2.46 | 0.65 | 2.26 | 0.78 | 1.99 | 0.37 | 1.74 | 0.41 |
| Percentage of significant fits (Lag-Opt) p<0.05 | Negative | 16.32 | 3.21 | 13.35 | 2.50 | 14.52 | 3.14 | 14.66 | 3.92 | 15.14 | 5.29 |
| Percentage of significant fits (Lag-Opt) p<0.05, Šidák | Negative | 4.50 | 2.10 | 2.28 | 1.13 | 0.82 | 0.19 | 1.05 | 0.67 | 0.85 | 0.88 |

**Supplementary Table 2.** Means and standard deviations (Stdev) across the 9 subjects for parameters summarized across GM voxels. Median CVR values that were taken from statistically thresholded distributions also did not include voxels where lag was found at the boundary condition.
